# Regulation of epigenetics and chromosome structure by human ORC2

**DOI:** 10.1101/2024.12.18.629220

**Authors:** Zhangli Su, Mengxue Tian, Etsuko Shibata, Yoshiyuki Shibata, Tianyi Yang, Zhenjia Wang, Fulai Jin, Chongzhi Zang, Anindya Dutta

## Abstract

We report a multi-omics study in a human cell line with mutations in three subunits of Origin Recognition Complex (ORC). The ORC subunits bind to DNA independent of each other in addition to as part of a common six-subunit ORC. DNA-bound ORC2 compacts chromatin and attracts repressive histone marks to focal areas of the genome, but ORC2 also activates chromatin at many sites and protects the genes from repressive marks. The epigenetic changes regulate hundreds of genes, including some epigenetic regulators, adding an indirect mechanism by which ORC2 regulates epigenetics without local binding. DNA-bound ORC2 also prevents the acquisition of CTCF at focal sites in the genome to regulate chromatin loops and indirectly affect epigenetics. Thus, our study reveals the genes and ORC1 regions bound by individual ORC subunits and suggests their role as epigenetics and chromosome structure regulators, independent of the role of the six-subunit ORC in DNA replication.

## INTRODUCTION

The Origin Recognition Complex, ORC, is a six-subunit complex that was identified in *S. cerevisiae* because of its essential role in DNA replication initiation. Five of the six subunits in yeast are AAA+ ATPases that associate through their C-terminal WH domains to form an open washer-shaped complex which binds DNA and bends the same, but in humans only the ORC1:ORC4 interface is an active ATPase (Bell and Stillman, 1992; Bleichert et al., 2015; Clarey et al., 2006; Dhar et al., 2001; Neuwald et al., 1999). Independent of its role in DNA replication, ORC has also been implicated in regulating chromatin and chromosome structure. Very early after its discovery, yeast ORC was implicated in silencing the HMR and HML loci in *S. cerevisiae* (Bell et al., 1993; Foss et al., 1993; Micklem et al., 1993). This mechanism involves ORC binding to those loci followed by the recruitment of SIR1 of the SIR1-4 complex, through the interaction of the BAH domain of ORC1 with SIR1 (Callebaut et al., 1999; Fox et al., 1997; Triolo and Sternglanz, 1996; Zhang et al., 2002). Separation of function mutations in ORC5 indicated that the replication initiation function of ORC can be separated from the transcription silencing function (Dillin and Rine, 1997; Fox et al., 1995). Furthermore, Drosophila ORC2 could complement the silencing defect of yeast *orc2-1*, while failing to complement the replication initiation defect, suggesting both that Drosophila ORC has silencing activity and that the silencing and replication initiation functions of ORC can be separated (Ehrenhofer-Murray et al., 1995). Indeed, Drosophila ORC, is also important for maintenance of the heterochromatin state through the ability of ORC1 to associate with and recruit HP1, a protein important for spreading and maintaining the heterochromatin state marked by the H3K9me3 chromatin mark (Cloos et al., 2006; Lachner et al., 2001; Pak et al., 1997).

In mammalian cells, the effect of ORC on chromatin structure has been less clear. Human ORC1, ORC2 and ORC3 associate with heterochromatin, specifically with HP1a and HP1b, and acute depletion of ORC subunits lead to loss of HP1 proteins from centromeric heterochromatin. ORC and HP1 have been suggested to together regulate the higher order heterochromatin structure at centromeric repeats (Prasanth et al., 2004; Prasanth et al., 2010). HP1 is known to associate with H3K9me3 modified chromatin to stabilize the repressive state in heterochromatin loci (Machida et al., 2018). ORC interacts with nucleosomes decorated by the repressive marks, H3K9me3, H3K27me3 and H4K20me2/3, and at least part of this association is mediated by an ORC associated factor, ORCA or LRWD1 (Bartke et al., 2010; Lukauskas et al., 2024; Shen et al., 2010; Vermeulen et al., 2010). The association of ORCA/LRWD1 with methylated histones has been proposed to be important for the heterochromatin structure (Chan and Zhang, 2012; Giri et al., 2015; Wang et al., 2017). However, deletion of ORCA/LRWD1 did not prevent the association of ORC with centromeric heterochromatin, and there was very little change in transcription of protein coding genes with a very slight de-repression of RNA from repetitive elements like satellite repeats, LINE and SINE elements (Kang et al., 2022). ORC1 has been shown to interact with Retinoblastoma (RB), H3K9me3 writer SUV39H1 and H3K9me3 to repress E2F1-regulated genes (Hossain and Stillman, 2016). In addition, acute depletion of ORC2 by siRNA resulted in chromosome compaction defects, leading to the appearance of “thick dumpy” chromosomes in 80% of metaphase spreads, suggesting defects in higher order structures of chromosomes (Prasanth et al., 2004). However, human ORC binding sites are highly enriched in open chromatin and near transcriptional start sites (TSSs) (Miotto et al., 2016). Interestingly, humanized yeast ORC acquired similar preference to open chromatin (Lee et al., 2021).

Over the last few years, we have created a few human cancer cell lines by CRISPR-Cas9 mediated genome engineering where *ORC1*, *ORC2* and *ORC5* genes were engineered to have the ORFs disrupted. The relevant ORC subunit proteins cannot be detected by regular immunoblots and yet the cell lines survive, proliferate and replicate DNA with the normal complement of origins of replication (Shibata and Dutta, 2020; Shibata et al., 2016). Careful cell-cycle studies show that all three cell lines have a normal cell-cycle progression. In the *ORC2Δ* cell line a very minimal level (<0.25% of wild type levels) of a truncated protein can be detected that reacts with anti-ORC2 antibody and co-immunoprecipitates with ORC3 (Chou et al., 2021), and we have discussed this further in (Shibata et al., 2025). WT cells have about 150,000 molecules of ORC2, so that we have calculated that if this truncated protein is functional ORC2, there are only <350 molecules of the protein. Even if this residual truncated ORC2 protein exists at <350 molecules per cell in the *ORC2Δ* cells, it does not materially change the conclusions of this paper. Till now there is no evidence that *ORC1Δ* or *ORC5Δ* cells express any detectable ORC1 or ORC5. Thus, these cells provide an opportunity to carefully examine the role of ORC in regulation of epigenetics, gene expression and higher order chromosome structure as studied by Hi-C. Note that there is only minimal slowing of cell proliferation upon deletion of the individual ORC subunits, ORC2, ORC1 or ORC5 (Fig. 2A and 4A of Shibata et al., 2016; Fig. 3B of Shibata and Dutta, 2020). Similarly, there is no change in DNA replication in terms of duration of S phase (Fig. 2C and 4C of Shibata et al., 2016; Fig. 4 of Shibata and Dutta, 2020), origin selection (Shibata et al., 2025), inter-origin distance and fork progression rate (Shibata and Dutta, 2020; Shibata et al., 2016). Thus the epigenetic effects we report below are not affected by slowing of cell proliferation.

## RESULTS

### ORC2 deletion and rescue cell lines

*ORC2Δ* (ORC2B2) is a clonal cell line derived from HCT116 p53-/-, referred to as “Wild-type” or “WT”. A new clonal cell line (referred to as “*ORC2*-rescue”) was developed where ORC2 expression is restored to its wild-type expression level by stable over-expression of ORC2 in the *ORC2Δ* cells (**Figure S1A** and **S1B**). We have shown that the loss of ORC2 is accompanied by destabilization of its closely interacting partner, ORC3, and the complete absence of ORC2 and ORC3 on the chromatin fraction is accompanied by a drastic decrease in the chromatin-associated ORC1, ORC4 and ORC5 (Shibata and Dutta, 2020; Shibata et al., 2016). Thus, the *ORC2Δ* cells lost virtually the entire ORC from the chromatin, that are restored to chromatin when ORC2 is re-expressed (**Figure S1C**).

### ORC2 both prevents and acquires local H3K9me3 and H3K27me3 marks

Acute *ORC2* knockdown leads to the disruption of HP1 loci and destabilization of associated proteins ORC3 and ORCA (Prasanth et al., 2004; Prasanth et al., 2010; Shibata et al., 2016). Consistent with this, ORC2 and ORC3 chromatin association was decreased in *ORC2Δ* cells and restored upon ORC2 rescue (**Figure 1A**). Similarly, chromatin association of another ORC2 interactor, ORCA or LRWD1, was also affected (**Figure 1A**). However, global levels of HP1 and repressive histone modifications H3K9me1-3 and H3K27me3 were not decreased in the *ORC2Δ* (**Figure 1A-B**). Instead, an increase in HP1 and H3K9me was detected but not rescued by restoration of ORC2. Neither was there any change in the chromatin level of HBO1, a histone acetyltransferase, that is known to associate with ORC (Iizuka and Stillman, 1999).

**Figure 1.**
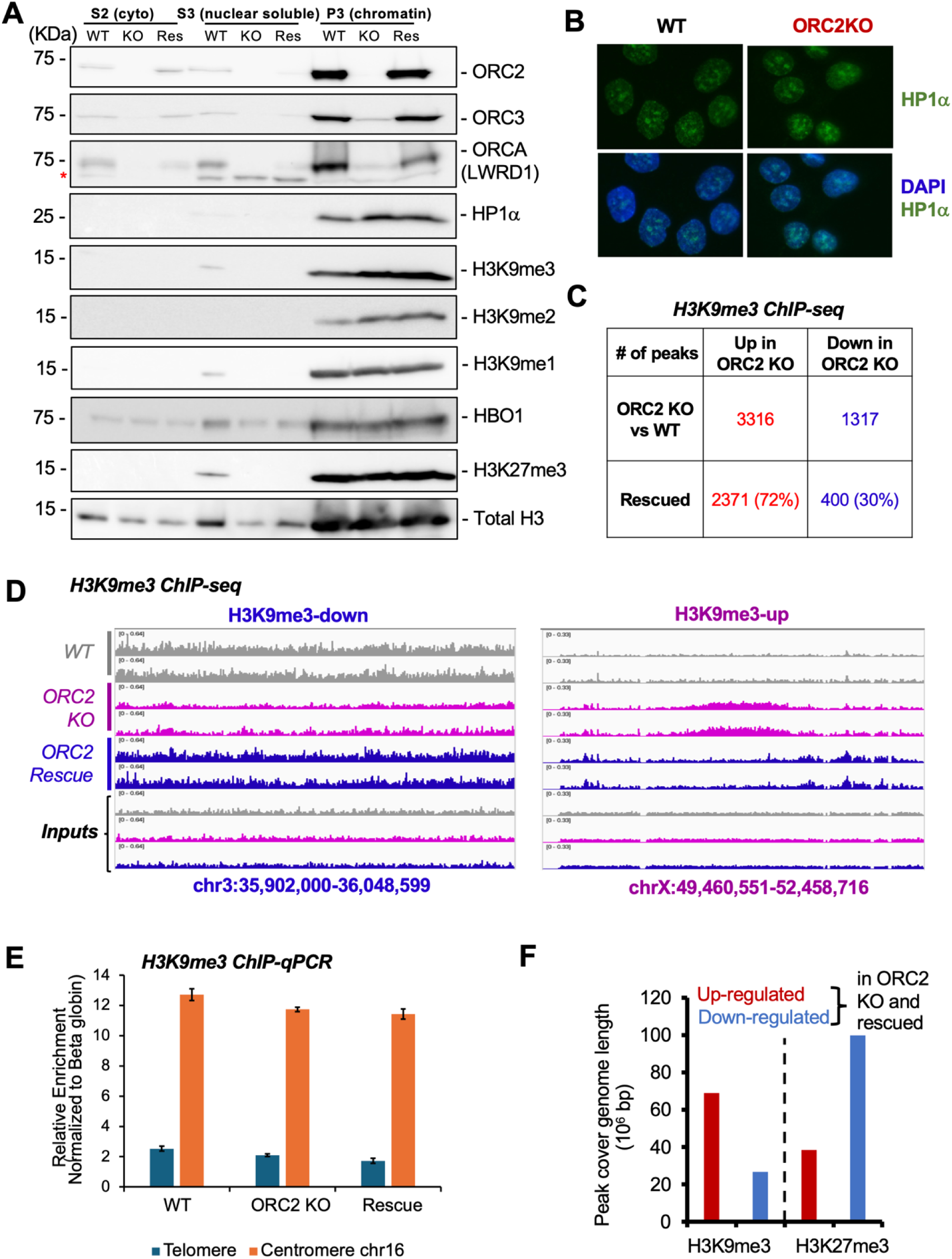
ORC2-dependent changes in H3K9me3 and H3K27me3 distribution. (A) Subcellular fractionation and western blotting of WT, ORC2KO and ORC2 rescue cells. S2 = cytoplasmic fraction, S3 = nuclear soluble fraction, P3= nuclear insoluble fraction. (B) HP1 foci are unchanged in ORC2KO cells. (C) H3K9me3 ChIP-seq peaks altered in knockout cells and rescued by reintroduction of ORC2. (D) Example H3K9me3 peaks that are dependent on ORC2, and down in the knockout (blue) or up in the knockout (purple) (n = 2). (E) ChIP qPCR shows that ORC2 knockout does not change H3K9me3 signal on telomeres or centromeres (n = 3). (F) Number of bases covered by ORC2 dependent H3K9me3 and K27me3. Red: upregulated in KO and rescued. Blue: downregulated in knockout and rescued.

Given ORC’s association with H3K9me3 and its role in HP1 recruitment, we anticipated its involvement in preserving H3K9me3 marks. Surprisingly, we observed more up-regulated H3K9me3 peaks than down-regulated peaks in *ORC2Δ* cells (**Figure 1C)**. We used SICER algorithm because it is optimized to identify broad histone modification islands. The 35,692 H3K9me3 peaks cover 800,593 kb with median peak width of 5.8 kb, out of which increased 3,316 H3K9me3 peaks cover 68,969 kb in the genome, with a median peak width of 6.6 kb. Furthermore, the up-regulation of H3K9me3 was more frequently reversed by ORC2 rescue compared to the downregulation (**Figure 1C-D**). We consider the rescuable sites as the ones most reliably dependent on the presence of ORC2. This suggests that ORC2 actively prevents H3K9me3 establishment in specific genomic regions. While previous studies reported dysregulation of centromeric and telomeric sequences in ORC2 knockdown cells (Deng et al., 2007; Prasanth et al., 2004), our ChIP-qPCR analysis revealed no significant changes in H3K9me3 levels at these regions (**Figure 1E**).

We then profiled another heterochromatin mark H3K27me3, which has not been investigated in ORC mutant cells before. There were fewer H3K27me3 sites that were dependent on the presence of ORC2 (**Figure S1D**): 5362 sites were downregulated when ORC2 was knocked out, of which 623 sites (12%) were rescued when ORC2 was restored. In comparison, there were 3474 H3K27me3 sites that were up regulated in *ORC2Δ* and 984 of them (28%) were dependent on ORC2 because they were restored by ORC2 rescue. In several regions there is mutually exclusivity in being marked by H3K9me3 and H3K27me3 (**Figure S1E**). For example, in those regions where H3K9me3 increases in the knockout, there is a decrease of H3K27me3 in the same cells, and the reverse happens when ORC2 is restored. This reciprocity (anticorrelation of fold change) of the two signals at a given site (**Figure S1F-G**) makes it unlikely that ORC2 simply helps recruit the two epigenetic writers for H3K9me3 or H3K27me3 to specific parts of the genome.

Since H3K9me3 and H3K27me3 peaks tend to decorate large contiguous stretches of the genome, we also asked how many bases of the genome are covered by these two marks in an ORC2 dependent manner (rescued). Among the sites where the repressive mark is dependent on ORC2, H3K27me3 covers more of the genome than H3K9me3 (**Figure 1F**, blue bars), although there are the same unexpected parts of the genome where these repressive marks are blocked by the presence of ORC2 (up in the knockout, decreased by the rescue) (**Figure 1F**, red bars). Taken together, the peak counts or bases covered by the marks surprisingly show that while ORC2 is required for maintaining the two common repressive marks in parts of the genome, as expected from the previous Literature, it is equally important for blocking the same repressive marks over larger parts of the genome.

### ORC2-dependent chromatin repression is mediated by repressive H3K9me3 and H3K27me3 marks

To identify sites where the chromatin is actually compacted or decompacted by ORC2, we profiled chromatin accessibility by ATAC-seq in HCT116 WT, *ORC2Δ* and *ORC2*-rescue cell lines (**Figure S1A**). ATAC-seq showed nucleosomal DNA pattern in all three cell lines confirming the quality of the ATAC-seq libraries (**Figure S2A**). We found significant alterations in chromatin accessibility upon ORC2 depletion. Specifically, there are more regions with increased accessibility (11,206 peaks) than regions with decreased accessibility (3,640 peaks) (**Figure 2A**). Of the 11,206 sites with increased transposase accessibility in *ORC2Δ*, ∼9,278 were decreased by the restoration of ORC2, and we abbreviated these as “ATAC-up”) (**Figure 2B** and **2D**). On the other hand, ∼2,311 of the 3,655 regions with decreased accessibility in the *ORC2Δ* became accessible again by the restoration of ORC2 (abbreviated as “ATAC-down”) (**Figure 2C** and **2D**). The ATAC changes not reversed by ORC2 restoration are likely due to secondary changes during ORC2 manipulation and we decided to focus on the ATAC changes that are rescued by ORC2 restoration and are clearly dependent on the presence of ORC2 (ATAC-up or ATAC-down). The ATAC-up sites are more abundant than the ATAC-down sites (**Figure 2D**). Overall, all transposase accessible sites in WT cells are enriched in genes especially promoter and enhancer regions relative to the genome background as expected, but the sites regulated epigenetically by ORC2 (ATAC-up and ATAC-down sites) are even more enriched in enhancer regions compared to all ATAC sites (**Figure 2E**), suggesting specific regulation of enhancers by ORC2.

**Figure 2.**
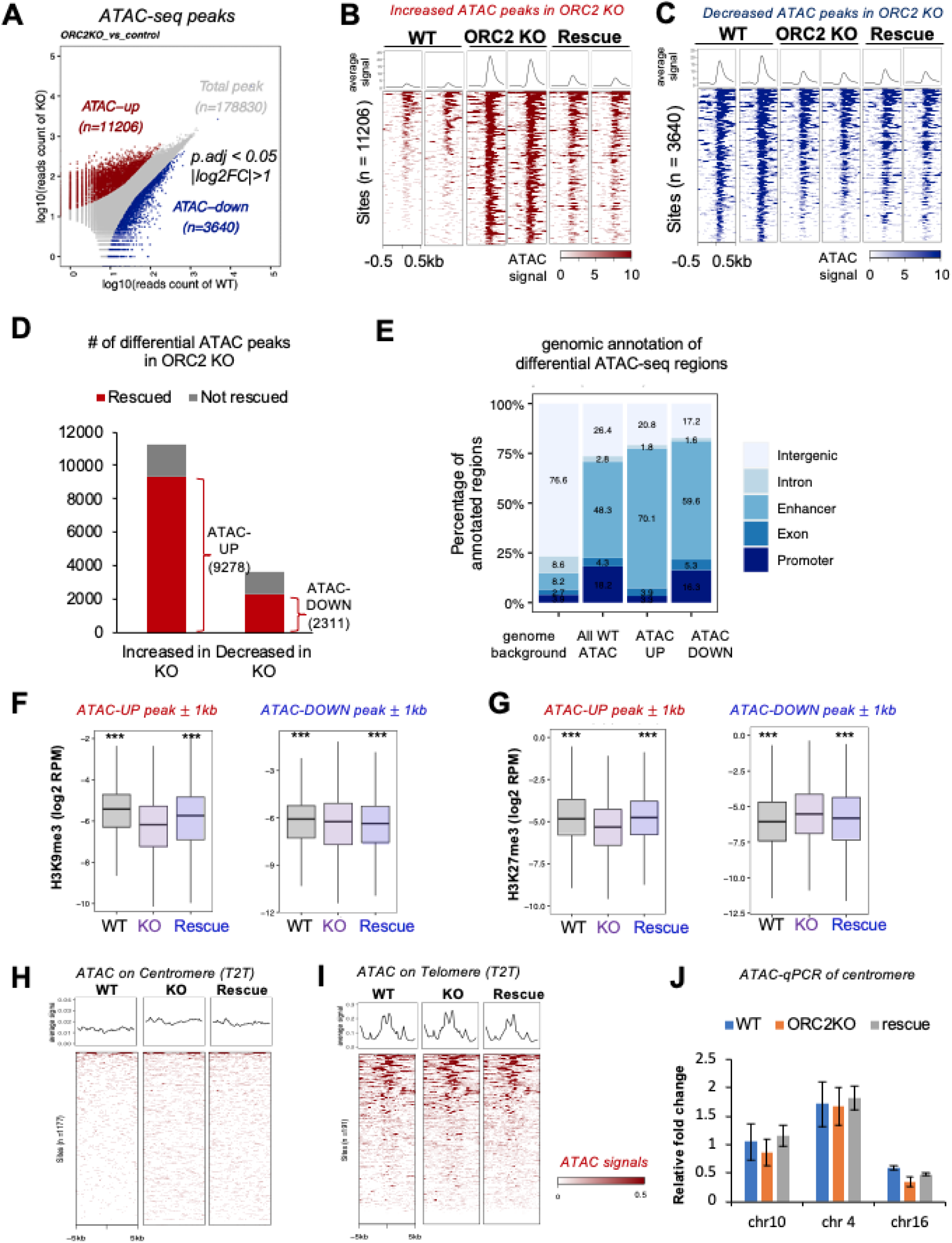
ORC2-dependent chromatin accessibility changes suggest its role in chromatin repression. (A) ATAC-seq peak signals compared in WT and ORC2KO cells. The up (red) and down (blue) peaks in ORC2KO are indicated. (B and C) Heatmap of ATAC signal in the three matched cell lines centered on peaks (B) upregulated or (C) downregulated in the KO cells (n = 2). (D) Number of ATAC peaks altered in the KO. Red: the sites where the changes were reversed re-introduction of ORC2. (E) Enrichment of ORC regulated ATAC peaks in enhancers. (F and G) (F) H3K9me3 and (G) H3K27me3 signal distribution on ATAC-up or ATAC-down peaks in three cell lines. *** p < 0.001 by student’s t test compared to the ORC2 KO. (H and I) Heatmap of ATAC signal centered on (H) centromeres and (I) telomeres in three cell lines. (J) qPCR of ATAC positive DNA at centromeres of three chromosomes.

A careful examination of the distribution of the H3K9me3 and H3K27me3 signal over the specific regions where transposase accessibility was dependent on ORC2 (ATAC-up and ATAC-down regions from **Figure 2D**) is shown in **Figure 2F-G and S2B-E**. The ATAC-up regions have significantly lower H3K9me3 and H3K27me3 signals in the *ORC2Δ* cells (**Figure 2F-G**) with a local dip (**Figure S2B-C)**, suggesting that some of the ORC2-mediated chromatin compaction in WT cells is accompanied by ORC2-dependent acquisition of these repressive marks. In contrast, the ATAC-down regions, where ORC2 prevents the compaction of chromatin, have a slight increase in H3K9me3 and H3K27me3 in the KO cells (**Figure 2F-G and S2D-E**), but the heatmaps suggest the increase is over the region and not specifically only at the sites.

Globally, centromeric accessibility was slightly increased by ORC2 depletion, consistent with previous results that ORC compacts centromere, but this was not rescued (**Figure 2H**). qPCR validation did not confirm the increased accessibility at three centromeres in *ORC2Δ* cells (**Figure 2J**). Telomeric heterochromatin accessibility was unaltered by ORC2 depletion and restoration (**Figure 2I**). Thus, although ORC regulates chromatin accessibility at many sites in the genome, we do not see any changes on centromeric and telomeric chromatin.

### ORC2-dependent chromatin accessibility changes are accompanied by changes in gene expression

To investigate whether the chromatin accessibility changes are accompanied by gene expression changes, we performed RNA-seq in the three cell lines (WT, *ORC2Δ* and *ORC2*-rescue). 1250 genes showed a significant increase and 541 genes showed a significant decrease in RNA levels by ORC2 depletion (**Figure 3A**). The increased RNA expression was reversed in 66% of the 1250 genes when ORC2 was restored (**Figure 3B-C**), confirming that ORC2 represses these genes; whereas the decreased RNA expression was reversed in a smaller percentage of gene after restoration of ORC2 (**Figure 3B** and **3D**). This pattern is reminiscent of the ATAC changes (**Figure 2**) in that more of the genes activated in *ORC2Δ*, than repressed in *ORC2Δ*, are reversibly linked to ORC2 levels. The magnitude and direction of all epigenetic changes are significantly correlated with the corresponding RNA expression changes in the *ORC2Δ* cells (**Figure 3E**).

**Figure 3.**
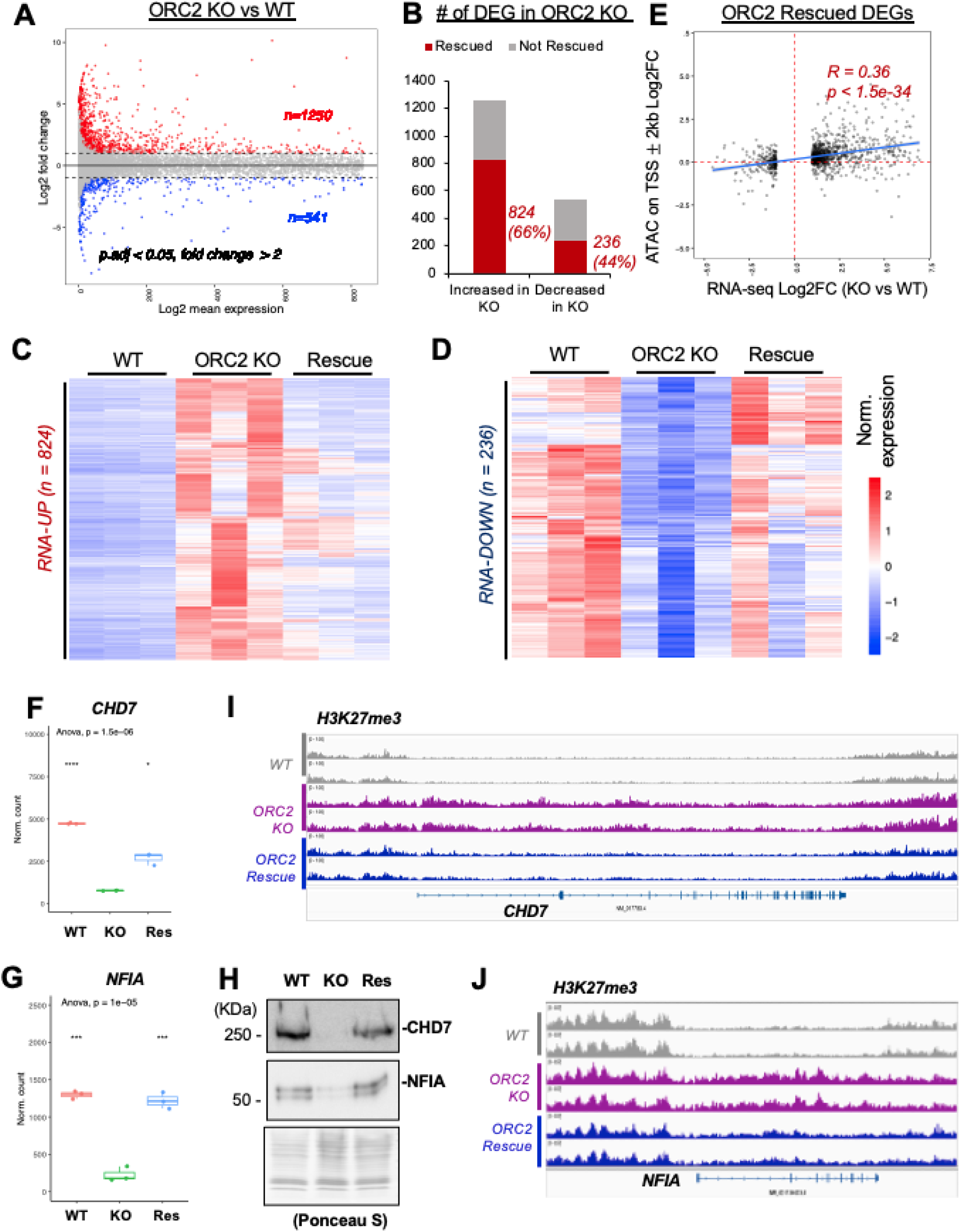
ORC2-dependent chromatin accessibility changes are accompanied by changes in gene expression. (A) RNAseq signal of different genes plotted to show mean expression and fold change in ORC2 KO cells relative to WT (n = 3). Genes significantly up-regulated (red) and down-regulated (blue) in the KO cells. (B) Number of genes whose expression is altered in the ORC2 KO. Red: genes whose expression change is reversed upon re-introduction of ORC2. (C-D) Heatmap of expression levels of genes that are (C) up-regulated or (D) down-regulated in the KO and rescued (n = 3). (E) Expression changes of genes in ORC2 KO cells correlated with change of ATAC accessibility in the TSS region. (F-G) CHD7 and NFIA RNA downregulated in an ORC2 dependent manner (n = 3). Global comparison was by Anova test across three cell lines, and pair-wise comparison with ORC2 KO was done with student’s t test (*** p < 0.001, * p < 0.05). (H) Western blot shows CHD7 and NFIA protein decrease on chromatin in ORC2 KO and rescue by ORC2. (I-J) H3K27me3 ChIP-seq signal over the CHD7 and NFIA gene in the three cell lines (n = 2).

To determine whether there were specific cellular pathways that were regulated at the gene expression levels, we carried out gene set enrichment analysis (**Figure S3A**). Pathways related to the specification of the extracellular matrix (ECM) and components involved in the interaction of the cell with the ECM are increased in the *ORC2Δ* relative to the WT cells. Conversely, the same pathways are repressed by the re-introduction of ORC2 into the KO cells (**Figure S3B**). Thus, ORC2 has a role in suppressing the extracellular environment and interactions that are associated with differentiated epithelial cells. Genes in pathways related to DNA and chromatin regulation (helicases, ATP dependent activity acting on DNA, binding to DNA secondary structure or to single-stranded DNA and nucleosome and histone binding) are repressed in the *ORC2Δ* cells (relative to WT) and induced in the rescued cells (relative to the *ORC2Δ*) (**Figure S3A-B**). Interestingly, genes involved in origin binding during DNA replication initiation are upregulated when ORC2 is restored to the *ORC2Δ* cells. This implies that gene expression that is dependent on ORC2 could be an integral component of cell-cycle progression, particularly DNA replication. Note, however, that the *ORC2Δ* cells do not show a significant change in cell cycle phase distribution or progression through S phase (Shibata et al., 2016), suggesting that change in replication factor or ATPase expression is subtle and does not impact cell cycle progression.

We have listed the genes whose changes in expression were definitely regulated by the presence or absence of ORC2 (**Table S1**).The ORC2-dependent changes in chromatin factors CHD7 (chromodomain helicase DNA binding protein 7) and NFIA (nuclear factor I A) were followed up as examples of factors that may indirectly cause some of the ORC2 dependent changes in gene expression and chromatin changes at sites where ORC2 does not bind (Basson and van Ravenswaaij-Arts, 2015). CHD7 and NFIA RNA and proteins both show ORC2-dependent expression (**Figure 3F-H**). The increase of H3K27me3 over CHD7 and NFIA locus in *ORC2Δ* cells could explain this regulation (**Figure 3I-J**).

### ORC2-dependent chromatin accessibility and RNA expression changes are specific to ORC2 and not ORC1 or ORC5

Since ORC2 is part of the six-subunit ORC complex, we wondered if other ORC subunits ORC1 and ORC5 have a similar phenotype in regulating chromatin accessibility and RNA expression (**Figure 4A**). To our surprise, loss of ORC1 and ORC5 (knockout cells generated in (Shibata and Dutta, 2020; Shibata et al., 2016)) showed different numbers of sites with accessibility changes compared to ORC2 (**Figure 4B, D** compared to **2A**). Consistent with the specific ATAC changes, we also observed specific gene expression changes in *ORC1Δ* and *ORC5Δ* compared to *ORC2Δ*. *ORC1Δ* has similar number of differentially expressed genes in both directions (522 up-regulated and 484 down-regulated) **(Figure 4C**), which correlated well with their ATAC changes in the TSS (**Figure S4A**). *ORC5Δ* has very minimal effect on chromatin accessibility and gene expression (**Figure 4D-E**).

**Figure 4.**
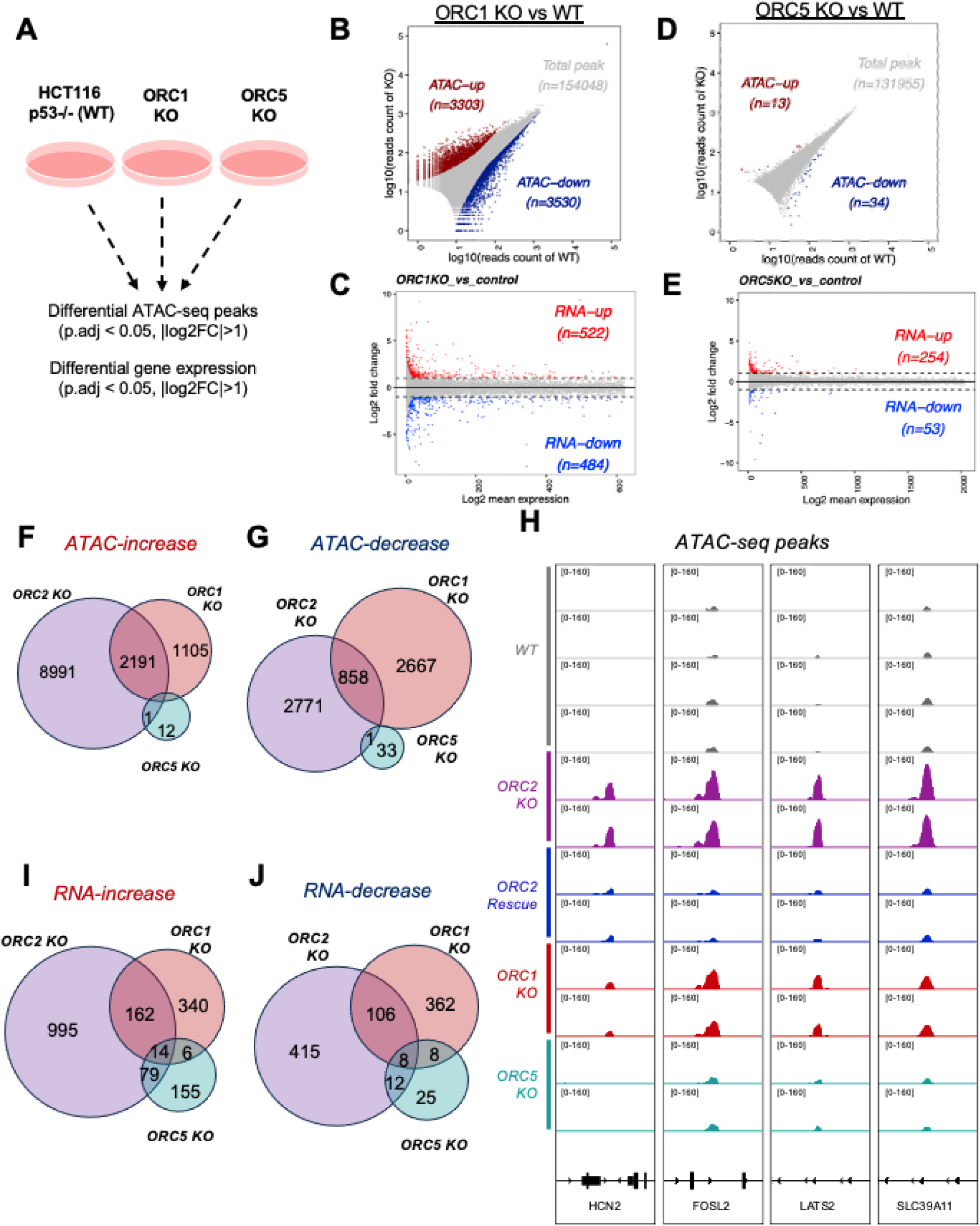
ORC2-dependent chromatin accessibility and gene expression changes are specific to ORC2 and not ORC1 or ORC5. (A) Schematic of experiment. (B-C) Chromatin accessibility (ATAC-seq) and gene expression (RNA-seq) changes in ORC1KO. Signals significantly up-regulated (red) or down-regulated (blue) in the ORC1 KO cells. (D-E) Same as (B-C) except for ORC5KO cells relative to WT. (F-G) Venn diagram showing overlap of ATAC changes in the three KO cells. (H) Example ORC2-dependent ATAC changes (chr19: 608305-614483; chr2: 28407886-28411076; chr13: 21031256-21037283; chr17: 72,698,613-72,700,987). (I-J) Venn diagram showing overlap of gene expression changes in the three KO cells.

It is particularly striking that none of the ATAC-change and a very small minority of the RNA changes were common between the three deletion lines (**Figure 4F-J**), as would have been expected if ORC regulated epigenetics in human cells as a holocomplex containing these subunits. There was more overlap in pair-wise comparisons. Up-regulated ATAC peaks in *ORC1Δ* have ∼66% overlap to the ATAC-up sites in *ORC2Δ*, while 24% of the down-regulated ATAC peaks in the *ORC1Δ* cells are also downregulated after *ORC2Δ* (**Figures 4F-G**). Examples of ORC2 ATAC-up sites (with ORC2-dependent chromatin closing) that are similarly regulated by ORC1, but not by ORC5, are shown (**Figure 4H**), as are examples of the larger number of ATAC-up sites that are regulated by ORC2, but not by ORC1 or ORC5 (**Figure S4B**). There are many more sites that are epigenetically regulated by ORC2 that are not regulated by ORC1 or ORC5. For example, 80% of the sites that depend on ORC2 for epigenetic repression are not dependent on ORC1 or ORC5 (**Figure 4F**), and 76% of the sites that depend on ORC2 for epigenetic activation are not dependent on ORC1 or ORC5 (**Figure 4G**). There are also sites that are epigenetically regulated by ORC1, but not ORC2 or ORC5 (examples in **Figure S4C**) or sites that are regulated by ORC5 alone (**Figure S4D**). Similarly, the overlap among the differentially expressed genes in each ORC subunit KO line is low with the highest number of genes uniquely dysregulated by *ORC2Δ* (**Figure 4I-J**), supporting the hypothesis of specific regulation by each ORC subunit. Although specific genes are dys-regulated in each KO line, some pathways are similarly altered in all three KOs, such as the up-regulation of “structural constituent of skin epidermis” (namely keratin genes) and down-regulation of “translation initiation factor binding” (**Figure S4E-F**).

### Binding sites of ORC1, ORC2 and ORC5

Historically ORC binding sites have been investigated by ChIP-seq for ORC1 or ORC2 proteins (Dellino et al., 2013; Long et al., 2020; Miotto et al., 2016; Tian et al., 2024). We decided to study ORC binding sites by a newer method, ChEC-seq because (a) it is not dependent on antibodies and access of antibodies to the epitope when a protein is bound in a complex to chromatin, and (b) our deletion cell lines allowed us to express the micrococcal nuclease bearing ORC subunit at the same level and without any competition from the endogenous ORC subunit (Zentner et al., 2015) (**Figure 5A**). The ORC subunit fused to micrococcal nuclease (MN-ORC1/2/5) was expressed from a doxycycline inducible promoter in the corresponding KO cell line (**Figure S5A-C**). The doxycycline levels were titrated to ensure that the expression of MN-ORC is comparable to that of the endogenous ORC subunit in WT cells. When ORC2 is knocked out, ORC1-5 are decreased on the chromatin, but the MN-ORC2 restored the chromatin association of these subunits showing that the MN-ORC2 is functional (**Figure S5D**). The fused micrococcal nuclease cuts the DNA at sites around the binding site of the ORC subunit, releasing these DNA fragments for sequencing and mapping (**Figure 5A-B**). Note that the binding sites reported in **Figure 5B**, are the ones that reproduced in two independent clones of cells expressing the relevant MN-ORC subunit (**Figure S5E**).

**Figure 5.**
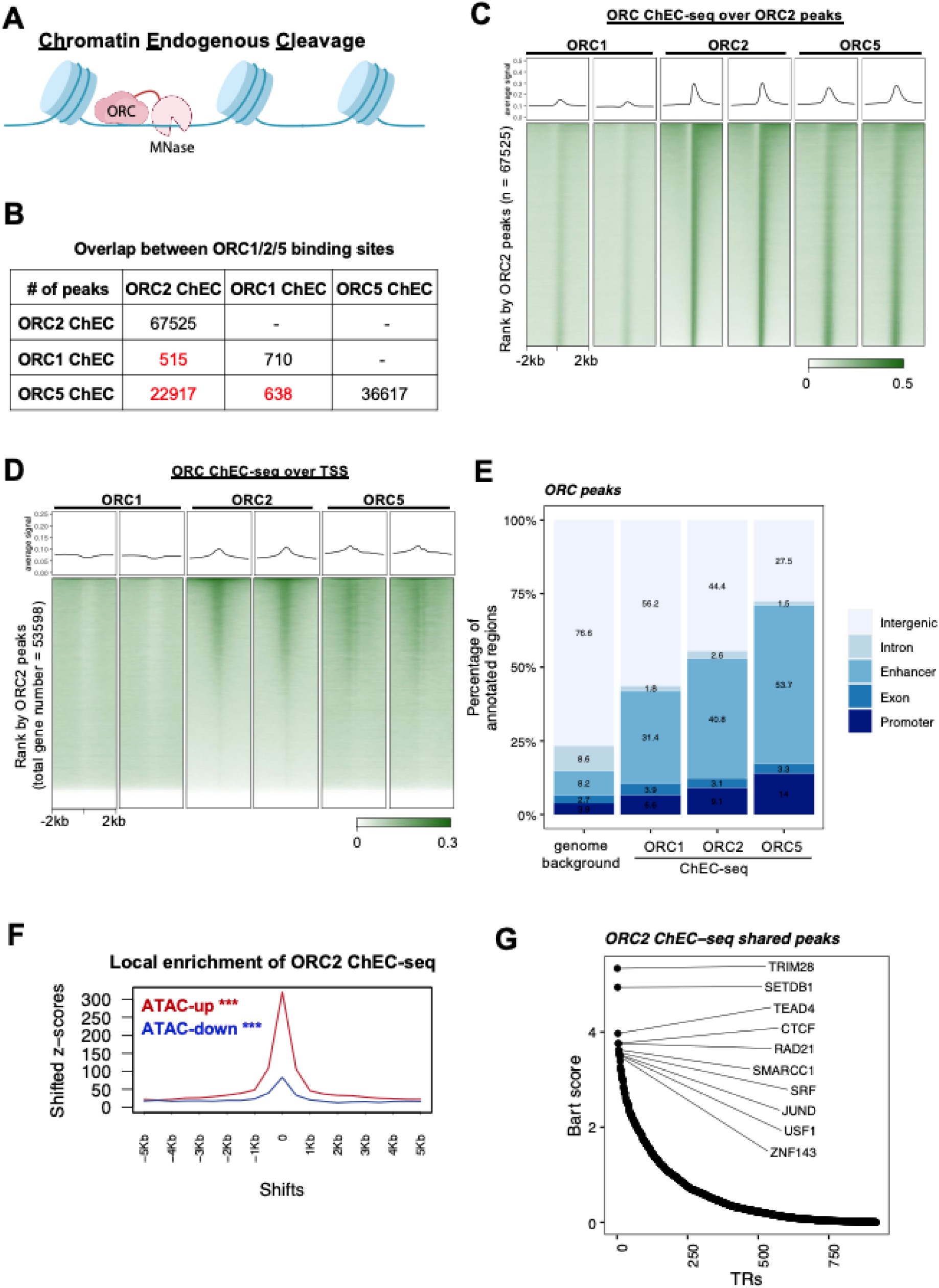
Binding sites of ORC1, ORC2 and ORC5. (A) Schematic of ChEC-seq experiments to identify binding sites of ORC subunits. (B) Number of peaks bound by each of the three ORC subunits and their overlap. (C-D) Heatmap of binding of the three subunits centered on sites bound by (C) ORC2 or (D) transcription start sites (n = 2). (E) Enrichment of binding sites of ORC subunits on enhancers and promoters. (F) ORC2 binding sites are enriched at sites that become more accessible (ATAC-up) when ORC2 is lost. A lesser enrichment is seen at ATAC-down sites. Z and p value is derived from permutation test (n = 1000), *** p < 0.001. (G) BART analysis transcription factors whose known binding sites are enriched near ORC2 binding sites.

The significant excess of ORC2 and ORC5 binding sites, relative to the ORC1 binding sites (**Figure S5F**) is consistent with previous observations, that the ORC2-3-4-5 complex is stable throughout the cell-cycle, while ORC1 is selectively degraded in S phase (Kara et al., 2015; Kreitz et al., 2001; Mendez et al., 2002; Tatsumi et al., 2003). Indeed previous ChIP-seq results showed that the number of ORC1 binding sites (120-4,000) is significantly lower than that of ORC2 binding sites (17,000-25,000, summarized in (Tian et al., 2024)). Also consistent with this is the observation, that ORC2 and ORC5 were bound to overlapping sites in the genome at approximately 22,000 sites, as would be expected if they bound DNA in the same complex. In contrast, ORC1 co-binding to ORC2 or ORC5 was restricted to ∼500 sites. To rule out that the low overlap of the ORC1 bound sites is not due to the poorer reproducibility of the ORC1 ChIP-seq (**Figure S5E**), we also counted the overlap of the union of ORC1 binding sites (rather than the intersection) with union ORC2 and union ORC5 binding sites and found the same result (**Figure S5P**). Consistent with this, heat maps of ORC1-, ORC2- and ORC5-binding across sites where ORC2 peaks were called, show that ORC2 and ORC5 have strong peaks at these sites, but although one can see ORC1 binding many of the same sites, the binding intensity is very weak, explaining why the peak-calling software calls far fewer ORC1 peaks (**Figure 5C**). This is exactly what is expected if ORC1 binds to subsets of ORC2-5 complexes in different cells in the population. We see the same trend in heatmaps of subunit binding across transcription start sites (**Figure 5D**). Taking both the called peaks and the heat maps into account, we conclude that ORC2 and ORC5 bind to tens of thousands of sites in the genome, both together and independent of each other and also independent of ORC1. This result is consistent with the observations in **Figure 3** and **4** that ORC2, ORC1 and ORC5 regulate the epigenetic state and expression levels of genes often independent of each other. In other words, the individual ORC subunits bind to DNA and carry out gene regulatory functions independent of the entire ORC holocomplex. Examples of sites with differential binding by ORC subunits and the ATAC changes when the ORC2 subunit is knocked out are shown in **Figure S5G**.

Even though the individual ORC subunits bound to many sites separate from each other, all the binding sites were enriched in enhancers and promoters relative to other parts of the genome (**Figure 5E**). The increase of ORC2 and ORC5, but not ORC1, at transcription start sites (promoters) is also evident (**Figure 5D**). The ORC2 ChEC-seq sites were highly enriched relative to random expectation to known ORC2 ChIP-seq sites, confirming that ORC2Chec-seq identifies *bona fide* ORC2 binding sites (**Figure S5H**). ORC2 bound sites are also enriched in areas with open chromatin (Transposase accessible ATAC sites) and sites with activated chromatin decorated by, H3K4me1and me2, H2AZ, H3K9Ac and H3K27Ac (**Figure S5H**). Although there was some enrichment relative to random expectation with sites carrying the repressive marks, H3K9me3 and H3K27me3, this enrichment was much less than with the activating marks. Overall, these results are consistent with the idea that ORC2 binds to open chromatin in WT cells and, based on the repression at many sites in *ORC2Δ* cells, has an active role in keeping the chromatin accessible.

We next analyzed whether ORC2 binding to DNA was correlated with the epigenetic repression by ORC2. There was a strong enrichment of ORC2 binding sites near ATAC-up sites, sites that are normally repressed by ORC2 and so become more accessible to transposase when ORC2 is knocked out (**Figure 5F**). Indeed, 5,597 (∼50%) of the ATAC-up sites overlapped with ORC2 binding sites (**Figure S5I**). In contrast, there was lower enrichment of ORC2 binding sites near the ATAC-down sites (**Figure 5F**) and only 785 (22%) of ATAC-down sites overlapped with ORC2 binding sites (**Figure S5J**). This is consistent with the rescue experiments, where the opening of chromatin in ATAC-up sites were reversed in greater numbers and more significantly when ORC2 was reintroduced in the KO cells (**Figure 2**), than the ATAC-down sites. We next examined changes in H3K9me3 around ORC2 binding sites in the *ORC2Δ* cells. Sites where H3K9me3 was down-regulated in *ORC2Δ* cells (“H3K9me3-down”) were enriched centered on ORC2 binding sites, and conversely H3K9me3-up regions were dis-enriched around ORC2 binding sites (**Figure S5K**). This is consistent with the hypothesis that ORC2 promotes or stabilizes H3K9me3 at many sites where it binds. Thus, repression of chromatin is correlated with ORC2 binding to many sites on the DNA. Finally, we analyzed the distribution of ORC2 binding sites at or near genes that were down- or up-regulated when ORC2 was deleted and rescued (**Figure S5N-O**). 60-70% of the genes regulated by ORC2, have ORC2 (and 5) binding sites in their gene body, and so are likely directly regulated by ORC2 (**Figure S5N**). This also suggests that 30-40% of the RNA-seq changes are indirect effects of ORC2, for example by regulating other transcription factors like CHD7 and NFIA and several others that we mention in the manuscript. The distribution of the ORC binding sites on the genes are shown (**Figure S5O**). For both sets of genes, RNA-down or RNA-up in ORC2 KO, ORC2/5 are enriched near TSS, suggesting that they could have either activation or silencing functions on different sets of genes, recruiting transcription silencers or activators depending on the gene context.

BART analysis determines whether any known factor binding sites are enriched among the sites where the ORC subunits bind. ORC2 binding sties were enriched in experimentally determined binding sites of chromosome regulatory factors including TRIM28, SETDB1, TEAD4, CTCF and RAD21 (the latter a part of the cohesin complex) (**Figure 5G**). Consistent with the observation that ORC1 and ORC5 bound to many sites in the genome independent of ORC2, BART analysis reveals that their binding sites are enriched in different subsets of chromatin binding proteins (**Figure S5L-M**). There are a few common hits, particularly the appearance of CTCF and RAD21 among the top hits for both ORC2 and ORC5, consistent with there being ∼22,000 sites in the genome co-bound by these two ORC subunits. Since both CTCF and cohesin are important for chromosome looping and establishment of chromosome domains, this observation led us to examine changes in chromosome 3D structure when ORC2 was deleted.

### ORC2 is required for specification of some chromosome compartments and some chromatin loops

ORC2 knockdown caused a change in the morphology of mitotic chromosomes (Prasanth et al., 2004), suggesting that ORC2 has a role in specifying 3-D chromosome structure. The changes in heterochromatin marks and chromatin accessibility seen with ORC2 deletion and rescue (**Figure 1-2**), also suggests that there may be accompanying changes in chromatin compartmentalization or looping. To this end, we generated high quality Hi-C data, as evidenced by the large number of cis- and trans-interactions (**Figure S6A**).

The bulk of the chromosome compartments are unchanged by deletion of ORC2, or the restoration of ORC2 (**Figure S6B**). Of the 12,397 chromosome compartments seen in WT cells (7,374 compartment A and 4,103 compartment B), there were 740 A-to-B and 104 B-to-A changes in ORC2 KO. Sites with B (closed, inactive expression) to A (open, active expression) compartment changes show local enrichment of ORC2 binding, whereas sites with A to B changes show local dis-enrichment of ORC2 binding (**Figure S6C**), indicative of ORC2 having a role in keeping adjoining areas closed in the WT cells. This is consistent with the ORC2 binding sites being enriched in sites where H3K9me3 is downregulated in the *ORC2Δ* cells (H3K9me3-down in **Figure S5I**). 15 stretches of chromatin changed upon ORC2 deletion from compartment B (closed) to A and returned to compartment B when ORC2 was restored. 12 stretches of chromatin changed in the opposite direction (from compartment B to A) and were rescued by ORC2. An example of ORC2-dependent compartment on chromosome 16 with the corresponding H3K9me3/27me3 changes is shown in **Figure 6A**. Note this change is only over ∼250 kb and the overall A/B compartment pattern is largely unaltered on chromosome 16 (**Figure S6D**). Change in compartment specification requires the PC1 value to change sign from plus to minus and vice-versa, and so may under-represent the interactivity changes when ORC2 is deleted and restored. Indeed when we take PC1 value as a continuous variable that measures interactivity, the difference in PC1 between the WT and the KO is significantly correlated with the difference between the Rescue and the KO (**Figure 6B**). Thus, globally there are many subtle ORC2 dependent interactivity changes, but these are not large enough to change the sign of the PC1 and thus change the compartment specification.

**Figure 6.**
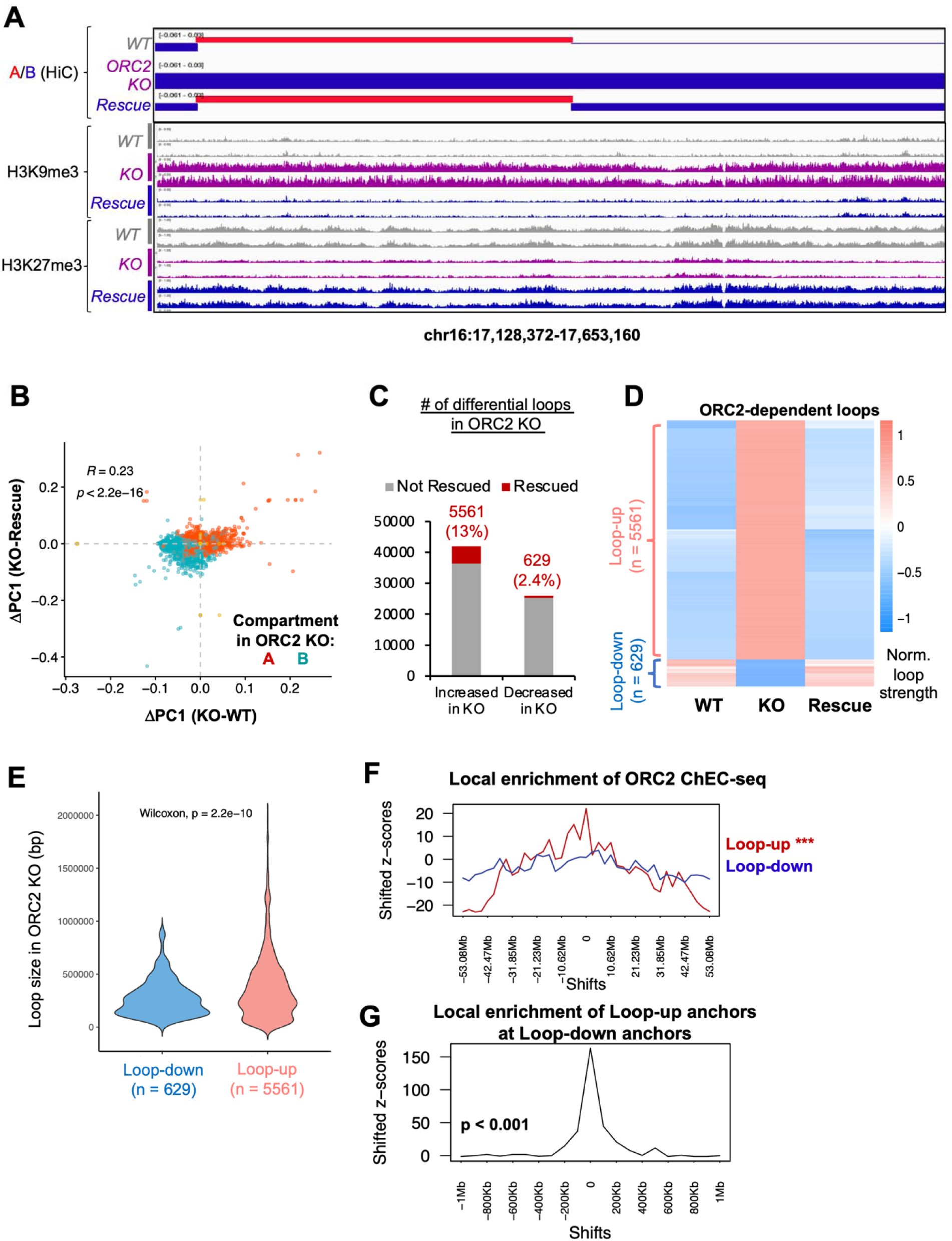
ORC2 is required for specification of some chromosome compartments and some chromatin loops. (A) Example compartment changes on Chr 16 from A (red) to B in the ORC2KO cells and is rescued by ORC2 restoration. The H3K9me3 and H3K27me3 tracks (n = 2) are shown. (B) Changes in PC1 when the KO is compared to the WT (X-axis) or the rescue (Y-axis) are correlated with each other at many more sites than show sign (compartment) changes. Color of the dots represent compartment status in the ORC2 KO (red: A, cyan: B). (C) Number of loops that increase or decrease in the KO relative to WT, and the portion that is rescued by ORC2 restoration. (D) Heatmap of loop strength of the loops (up or down) that are rescued in (C). (E) Loops that become stronger in the KO are longer than the loops that become weaker. (F) Enrichment of ORC2 binding sites across the region centered on loop-up or loop-down sites. (G) Enrichment of loop-up anchors across the region centered on loop-down anchors.

The Hi-C analyses also revealed that although there were tens of thousands of loops that were increased or decreased in the KO or the rescued cell lines (**Figures S6E**) the formation or suppression of a more limited set of chromatin loops was dependent purely on whether ORC2 was present or not. 5561 loops were increased in the *ORC2Δ*, and re-suppressed when ORC2 was restored (“Loop-up”) (**Figure 6C-D**). In contrast, only 529 loops were decreased in the *ORC2Δ* and restored when ORC2 was restored (“Loop-down”). The lengths of the Loop-up loops were greater than the lengths of Loop-down loops (**Figure 6E**), suggesting there are more long-range interactions in the *ORC2Δ*. There is significant local enrichment of ORC2 binding sites at Loop-up but not Loop-down regions (**Figure 6F**). There is also a significant overlap between the ATAC-up sites and Loop-up sites, suggesting that acquisition of new loops in the KO cells is accompanied by local opening of the chromatin, which may itself be due to the loss of ORC2 in those areas of the chromosomes (**Figure S6F**). Interestingly, the loop-up anchors are strongly enriched near the loop-down anchors (**Figure 6G**), suggesting ORC2 re-organizes loops using the same anchors.

### A functional interaction between ORC2 and CTCF driven by local accessibility

Because CTCF bound sites often anchor chromatin loops and because ORC2 binding sites were enriched in CTCF binding sites (**Figure 5G**), we wondered whether CTCF binding to the chromatin is directly or indirectly regulated by ORC2 binding to local sites, thus explaining why some of the loops were dependent on ORC2. We carried out cut-and-run to identify the CTCF binding sites in the WT, *ORC2Δ* and ORC2-rescue cells (**Figure 7A**). ∼14,000 CTCF bound sites were unchanged by the changes in ORC2 in the three cell lines. There were, however, ∼9,600 sites where CTCF binding increased in the *ORC2Δ* cells (CTCF-up sites), and which were decreased when ORC2 was restored. In contrast, CTCF binding was decreased in only ∼360 sites in the KO cells in an ORC2 dependent manner (CTCF-down sites).

**Figure 7.**
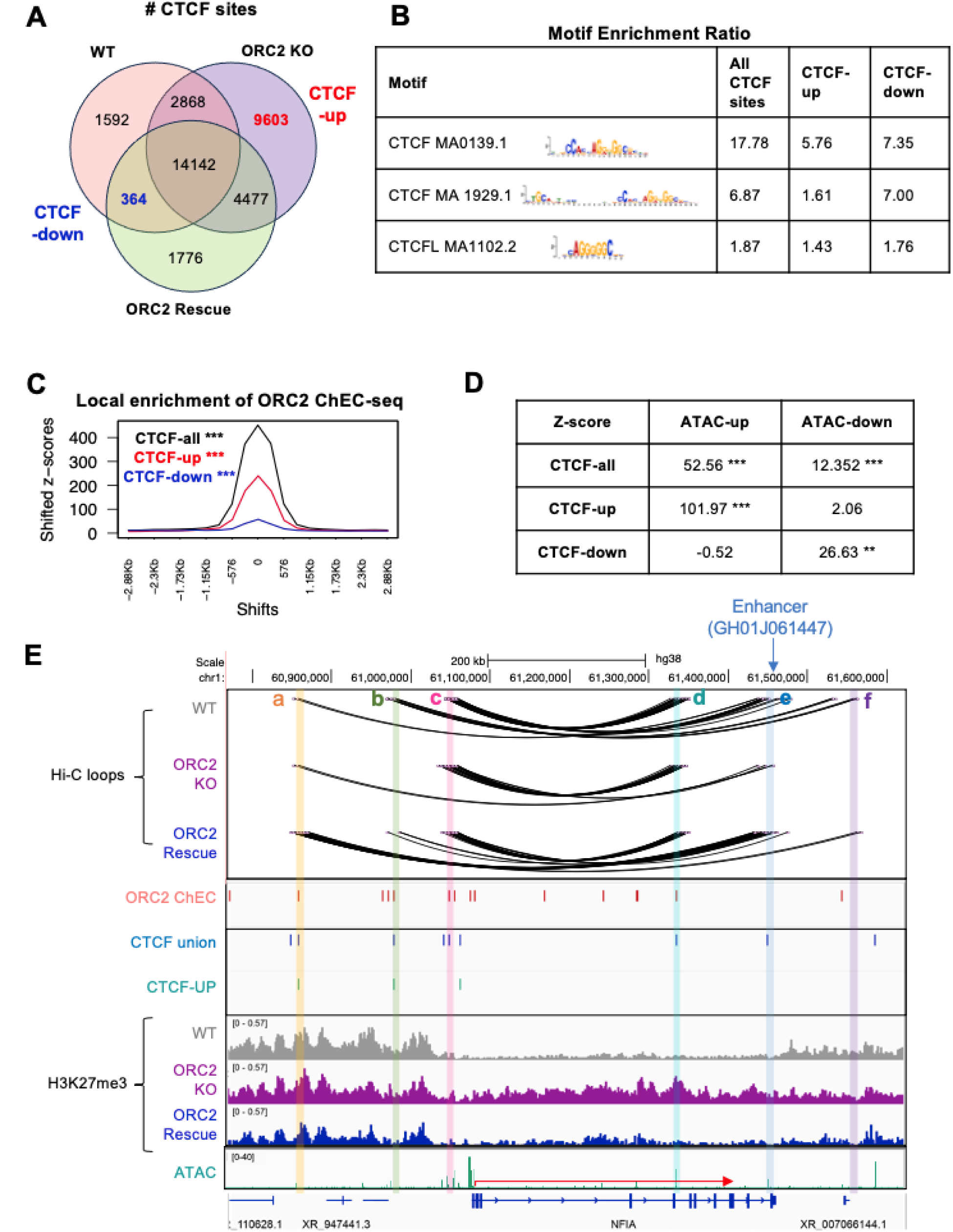
A functional interaction between ORC2 and CTCF driven by local accessibility. (A) Venn diagram of CTCF binding sites in the three cell lines. Red: CTCF binding sites that are increased in the KO and decreased in the rescue. Blue: CTCF binding sites present in the WT that disappeared in the KO and re-appeared when ORC2 is restored. (B) CTCF motif enrichment ratio in indicated classes of CTCF binding sites. (C) Enrichment of ORC2 binding sites across the region centered on CTCF-binding sites of the three classes by permutation test (n = 1000). (D) Enrichment of ATAC-up or -down sites at CTCF-binding sites of the three classes by permutation test (n = 1000). (E) Top three tracks: Loops in the NFIA locus in the three cell lines. The loop anchor sites a-f are indicated at the top and marked by the colored bars running through all tracks. ORC2 and CTCF binding sites are shown in the middle tracks. Bottom three tracks: ORC2 dependent prevention of H3K27me3 mark at the locus.

The CTCF-up sites have a lower CTCF motif enrichment ratio compared to all CTCF sites (**Figure 7B**) suggesting that these sites, not bound by CTCF in WT cells, have a lower affinity for CTCF. CTCF binding sites were among the top enriched sites near ORC2 bound DNA in BART analysis (**Figure 5G**), and 75% of union CTCF sites overlap with ORC2 ChEC-seq sites (**Figure S7A**). To determine whether ORC2 binding to DNA directly influenced CTCF binding to the same locations, we analyzed the overlap of ORC2 bound DNA (ORC2 ChEC-seq sites in the WT cells) with the CTCF-up or -down sites (**Figure 7C** and **S7B**). A very clear pattern emerged. While all CTCF bound sites were enriched in ORC2 bound sites, the CTCF-up sites showed a marked co-localization with ORC2 bound sites, compared to the behavior of the CTCF-down sites. In other words, many sites bound by ORC2, but not CTCF, in the WT cells, bound CTCF when ORC2 was lost (in the *ORC2Δ* cells), and this binding of CTCF was repressed when ORC2 was restored. 3,327 CTCF-up sites that were repressed by re-introduction of ORC2 overlapped with ORC2-bound sites. This is in contrast to only 99 of the CTCF-down sites overlapping with sites normally bound by ORC2 in WT cells.

ORC2 could repress CTCF binding by directly competing for CTCF binding sequences. However, motif-analysis of ORC2 ChEC-seq sites does not show any evidence that ORC2 binding sites have enrichment for CTCF motifs (data not shown). An alternative possibility is that ORC2 bound to DNA compacts chromatin (the ATAC-up sites in **Figure 5F**), and this prevents CTCF binding. Consistent with this, we see an enrichment of ATAC-up sequences with all CTCF bound sites, and particularly with CTCF-up sites **(Figure 7D**). An example of such a CTCF-up sites is shown near anchor point c of the NFIA locus in **Figure 7E**. In WT and ORC2 rescue cells enhancers near the 3’ end of the gene near anchor point e (GH0161447) interact with elements at anchor points b and c near the 5’ end of NFIA in WT and ORC2 rescue cells. We hypothesize that the appearance of a new CTCF site in the KO cells near anchor point c stops the extrusion of the chromatin loop and strengthens the c-d interaction, while decreasing the longer b-e loop that uses anchor-points further out. The weakened b-e interaction in the KO cells decreases the influence of the 3’ enhancer on the promoter of NFIA, and the resulting drop in transcription leads to the spread of H3K27me3 marks from the neighborhood on to the NFIA gene. Another example is shown in **Figure S7C** for CHD7, where a CTCF-up site is seen in anchor point c. In WT cells and ORC2 rescue cells, the dominant loop connects points a and e, bringing a distal enhancer GH08J060997 from e to the CHD7 promoter at a. The promoter could also be activated by several other enhancers in the a-e loop. In ORC2 KO cells, the new CTCF site near anchor point c stops loop exclusion of the c-e segment, so that the c-e loop strengthens at the expense of the outer b-e and a-e loops. The stronger c-e loop separates the CHD7 gene from the GH08J060759 enhancer and we propose that this insulation from the enhancer prevents CHD7 transcription and leads to H3K27me3 spread from neighboring regions into the CHD7 gene.

## DISCUSSION

### ORC subunits bind to many sites in the genome independent of each other

This multi-omics analyses of three cancer cell lines that have individually deleted three of the six subunits of ORC has yielded several unexpected observations. The most striking observation is that ORC subunits can bind to sites on the genome independent of other subunits and regulate the chromatin state independent of each other. Although ORC was discovered in yeast as a tight complex of six subunits, in human cells it was apparent that a core subcomplex of ORC2-3-4-5 was much more stable than the six subunit complex that also includes ORC1 and ORC6 (Dhar et al., 2001). Our earlier work had shown that even in the absence of ORC1, ORC2 or ORC5, ORC6 was still associated stably with the chromatin (Shibata and Dutta, 2020; Shibata et al., 2016). Also, we have published that deletion of ORC1 has no effect on chromatin loading of ORC2 and ORC5 (Fig. 3D, Shibata et al., 2016) and deletion of ORC5 has no effect on chromatin loading of ORC1 or ORC2 (Fig. 2E, Shibata and Dutta, 2020). In an earlier paper (Tian et al., 2024), we reported similar lack of overlap of published ORC1 and ORC2 binding sites from ChIP-seq experiments. As a positive control, we also looked at ChIP-seq data for other complexes like the SWI/SNF complex or the PRC2 complex and found a much higher overlap of binding sites between the subunits. These analyses collectively suggest that ORC subunits could bind to chromatin independent of each other. Now, from the ChEC seq results (**Figure 5**) we can say that although there are some sites where ORC2 and ORC5 co-bind to the genome stably, and although ORC1, ORC3, ORC4 and ORC5 require ORC2 for bulk association with chromatin (**Figure 1A and S1C**), there are many sites where ORC1, ORC2 and ORC5 bind independent of each other. The ATAC-seq and RNA-seq results (**Figure 4**) also support the idea that many more sites are regulated by the three subunits independently than as part of a holo-ORC that contains all three proteins.

The restoration of ORC2 (or ORC2-MNase in the *ORC2Δ* cells) stabilizes the ORC1 and ORC5 subunits on the chromatin, which seems to contradict the different binding sites of the subunits revealed by ChEC-seq. There are two possibilities: the stabilization happens in the nucleoplasm, but the stabilized subunits can bind to different cellular proteins and bind to different DNA sites in the genome independent of each other. Another possibility is that the bulk chromatin association of the ORC holocomplex is not being picked up in the ChEC-seq data, either because the MNase for all three subunits are being masked in the holocomplex (unlikely to happen with three subunits) or because the chromatin association seen in chromatin fractionation is too transient to be reported by ChEC-seq. The differences in ATAC-seq data between the three subunits can be more easily explained if different subunits regulate the local epigenetics differentially at various sites due to differences in the nearby co-factors.

### ORC-dependent epigenetic changes regulate gene expression

The gene expression changes produced by ORC2 deletion correlate well with the local changes in chromatin compaction, as measured by ATAC-seq. Here again, the activation of genes seen upon ORC2 loss is more markedly reversed when ORC2 is restored. Thus, the repression of genes by local, facultative, chromatin compaction seems to be more dynamically regulated by ORC2. Once again, the deletions in other subunits of ORC clearly show that different genes are dysregulated, so that each of the three ORC subunits appear to have their own independent spheres of influence. Gene-ontology studies suggest that the one common pathway upregulated by all three knockouts involves structural constituents of the skin epidermis, suggesting that ORC expression inhibits skin epidermal differentiation. On the other hand, DNA helicase pathway is repressed by ORC2 and ORC5 deletion, but up-regulated by ORC1 deletion. Protein folding chaperone binding pathway is down-regulated by ORC5 and up-regulated by ORC1. Collectively, these results underscore that the different subunits seem to regulate different sets of genes.

### Repression of chromatin by ORC

As cited in the introduction, studies in yeast, Drosophila and limited studies in humans showed that ORC was involved in repression of chromatin. The ATAC-seq and H3K9me3 and H3K27me3 ChIP-seq results support the fact that human ORC2 is involved in chromatin compaction (**Figure 1-2**), The enrichment of ORC2 binding sites with the ATAC-up sites (**Figure 5**) confirms that the binding of ORC2 to a local area facilitates chromatin compaction in WT cells. As discussed above, the ATAC-seq analyses also highlight that the three individual subunits studied regulate chromatin compaction independent of each other.

Examination of H3K9me3 and H3K27me3 marks in these three cell lines is complicated by the fact that these marks are often present over broad stretches of DNA, and not as sharp peaks like the ChEC-seq or ATAC-seq data. However, examining the enrichment of these marks over broad parts of the chromosomes revealed that H3K9me3 marks are downregulated around ORC2-bound sites when ORC2 is deleted (**Figure S5K**). Thus, it is likely that the chromosome compaction around ORC2 sites in WT cells is at least partly effected by the writing or stabilization of H3K9me3 marks. ORC has been proposed to help stabilize H3K9me3 marks and recruit HP1 to constitutive heterochromatic foci, but a role in maintaining H3K9me3 (or H3K27me3) marks in focal areas of euchromatin (facultative heterochromatin) has not been reported. Interaction of ORC2 with PRC2, the writer of H3K27me3 has been found by mass-spectrometry experiments (Oliviero et al., 2016), and so ORC2 could locally increase H3K27me3 by recruiting PRC2. ORC2 has been consistently pulled down by nucleosomes with H3K9me3/H3K27me3/H4K20me3 marks (Bartke et al., 2010; Lukauskas et al., 2024). ORC1 has been shown to interact with Retinoblastoma (RB), H3K9me3 writer SUV39H1 and H3K9me3 to repress E2F1-regulated genes (Hossain and Stillman, 2016). We have also summarized in **Supplementary Table 3** the interactions between epigenetic regressors and ORC2, ORC1 and ORC5 that are reported in BioGrid (ORC2 with LRWD1/ORCA, SIRT2, CBX3, CBX5, EZH2, LMNA; ORC1 with LRWD1/ORCA, ATRX, CBX1, DOT1L, PRC1, RB1; only LRWD1/ORCA with ORC5). Note that CBX3/CBX5 are part of the HP1 group, which binds to H3K9me3 and EZH2 is part of PRC2, responsible for H3K27 methylation.

Association of ORC binding sites with binding sites with repressive chromatin regulators is also suggested by the BART analysis (**Figure 5G**), which identifies known factor-binding sites that are enriched in the ORC subunit bound sites. For example, TRIM28 and SETDB1 (writer of H3K9me3) binding sites were enriched in the ORC2 and ORC1 binding sites. CBX3 (part of HP1) and ZNF274 are enriched near ORC1 binding sites. The TRIM28-SETDB1-ZNF274 complex is enriched near H3K9me3 sites (Frietze et al., 2010; Valle-Garcia et al., 2016), and HP1 is known to bind to H3K9me3 marks and stabilize them (Machida et al., 2018), and this may be one mechanism by which ORC2 promotes H3K9me3 marks and (with HP1) chromatin compaction in unique focal sites in the genome.

There have been several reports that ORC is involved in heterochromatinization of parts of the genome that have repeat sequences, e.g. pericentromeric satellite repeats, and this heterochromatinization is dependent on the acquisition of H3K9me3 marks (Decombe et al., 2021; Nishibuchi and Dejardin, 2017; Prasanth et al., 2010). However, our studies on the alpha satellites in the WT and the *ORC-subunitΔ* cells reveal that in these long-established cells lacking ORC subunits, the cells have compensated in some manner with no evidence of chromatin de-compaction or repeat RNA induction in the *ORCΔ* cells (**Figure 1E, 2H-J**, and data not shown). We reason such difference could be because the chronic depletion in our cell lines allows the centromeres to adapt and reacquire heterochromatin, while the previous reports acutely depleted ORC subunits. Similarly, ORC has been reported to localize to telomeres together with HP1 at telomeric loci (Deng et al., 2009). Again, in these cells where ORC has been stably deleted, we see no evidence of telomere decompaction, suggesting that although recruited to telomeres, ORC, does not have a direct role in telomere heterochromatinization.

### Activation of chromatin by ORC

The surprising result is that we also detected significant number of sites where ORC subunits seem to keep the chromatin open (ATAC-up sites). The ORC2 deletion shows that it keeps chromatin open by preventing the appearance of H3K9me3 and H3K27me3 marks. Here again, the enrichment of ORC2 binding sites with the ATAC-down sites (**Figure 5**) suggests that the binding of ORC2 to a local area facilitates chromatin decompaction in WT cells and several possible mechanisms are also suggested by our results.

Some of this activation could be mediated by chromatin activators that bind with ORC to these sites. BART analysis revealed the enrichment of DNA binding sites for SMARCC1 (an ATP dependent SWI/SNF type chromatin remodeler) and JUND (an AP1 transcriptional activator) with ORC2 and ORC5 binding sites. Therefore, these proteins may be involved in activating the chromatin at or near some ORC2 or ORC5 binding sites.

Another mode of activation of chromatin by ORC is through altering the regulatory loops on the chromosomes. ORC has been shown to co-localize with cohesin in Drosophila (MacAlpine et al., 2010). The enrichment of CTCF and RAD21 (a cohesin subunit) binding sites near ORC2 and ORC5 and the enrichment of SMC1A (also a cohesin subunit) binding site near ORC5, suggested that some of the human ORC subunits may be involved in chromatin looping. Consistent with this, our results suggest that ORC2 binding to the chromatin appears to keep CTCF away from certain sites, and this could be because of the local repression of chromatin at those sites. In the absence of ORC2, CTCF binds to these sites, creating new anchor points for loop formation, and thus increasing looping. In two loci studied, NFIA and CHD7, this increased looping isolates a known enhancer from the promoter of the gene, resulting in decrease in transcription and spread of repressive marks across the locus.

### Indirect regulation of epigenetic state by ORC

While ORC2 binding to DNA underlies some of the chromatin compaction in WT cells that is seen in ATAC-up sites, there are clearly many other sites where chromatin compaction is regulated by ORC2 (in either direction), but without any overlapping ORC2 bound sites. One major mechanism of this indirect regulation could be through ORC2 regulating chromatin remodeling factors and transcription factors. We have confirmed the changes in CHD7 and NFIA in this paper, but the list of genes whose transcription is dependent on ORC includes several other chromatin remodeling factors and transcription factors: YAP1, TCF7L2, KLF3, EGR3, EGR1, SCML1, HOXC13, ASXL2 and many others. Therefore regulation of the expression of CHD7, NFIA and these other factors could also dictate some of the ORC-dependent epigenetic changes that are not mediated by local binding of ORC.

### Chromosome compartmentalization and looping regulated by ORC

As with the ATAC-seq changes, most of the changes in chromatin compartmentalization and loops seen upon ORC2 deletion are not reversed when ORC2 is restored. However, the division of the chromosomes into compartments A and B uses a strict threshold, and so cannot identify sites where the restoration of ORC2 produced a partial restoration of the interaction frequency, but not sufficient to change the compartment denomination. Indeed this is supported when we simply look at the changes in interactivity, as measured by ΔPC1, in the ORC2 KO and rescue cells. Consistent with the observation that ORC2 mediated epigenetic suppression is most easily restored when ORC2 is restored (i.e. there are more ATAC-up sites), we see that the parts of the chromosomes that move from compartment B (compacted) to compartment A, are primarily the compartments that are reversed (back to compacted compartment B) when ORC2 is restored.

Focusing on the loops that are altered in a reversible manner, we find that ORC2 is required mostly for preventing excess loop formation. However, as with epigenetic changes that were mediated indirectly without local ORC2 binding, some of the ORC2-dependent loops are also being regulated indirectly by ORC2, not through direct binding of ORC2 near those loops, but through the regulation of other factors involved in loop formation. Interestingly, CHD7 knock-down (as seen in *ORC2Δ*) is recently reported to promote long-range interactions (Cheng et al., 2023).

In conclusion, this multi-omics analyses of five cell lines where levels of individual ORC subunits are manipulated provide a wealth of information about how ORC subunits regulate gene expression independent of their role as part of a replication initiator complex. More often than not, the three ORC subunits bind to chromatin and regulate epigenetic state and gene expression independent of each other rather than as part of a six-subunit holocomplex. Consistent with what was expected from the lower eukaryotes, we find thousands of sites where ORC subunits are required to facultatively repress chromatin and gene expression. However, a surprisingly large number of sites are regulated in the reverse direction such that ORC subunits keep chromatin open and facilitate gene expression. The genes regulated by individual ORC subunits are diverse and implicated in unexpected pathways like epithelial differentiation and are regulated by epigenetic changes mediated both by local binding of ORC subunits and indirectly by regulating the expression of other chromatin regulatory factors and transcription factors or the looping of chromosomes. ORC subunits have a role in protecting certain sites from binding by CTCF and thus preventing extraneous chromosome loop formation. This is the first detailed study of how and where ORC regulates epigenetics and gene expression in human cells and the unanticipated scale and breadth of the regulation opens new chapters in ORC biology.

## ACKNOWLEDGEMENT

This work was supported by NIH grants R01 CA60499 to AD, R35 GM133712 to CZ, and R00 CA259526 to ZS. We thank all members of both the Zang and Dutta Labs for many helpful suggestions.

## METHODS

### Cell lines and culture

HCT116 p53-/- cell (referred to as wild-type, RRID: CVCL_S744) was a generous gift from Fred Bunz (Johns Hopkins) (Bunz et al., 1998). ORC2B2 (referred to as ORC2-/-) was a clonal cell line that was established by CRISPR/Cas9 technology in the HCT116 p53-/- background (Shibata et al., 2016). ORC2 rescue cells were established by pLHCXORC2 cloning ORC2 with 3’UTR and 5’UTR and clonal selection. ORC1 or ORC5KO clones were established in HCT116 p53-/- cells using CRISPR/Cas9 technology (Shibata and Dutta, 2020; Shibata et al., 2016). All cell lines were maintained in McCoy’s 5A-modified medium (Corning, no. 10-050-CV) supplemented with 10% fetal bovine serum.

### ATAC-seq library preparation

ATAC-seq for cell lines was performed based on Omni-ATAC-seq protocol (Corces et al., 2017) with 50,000 cells. Nuclear pellet was subjected to transposition with Nextera DNA Sample Preparation Kit (Illumina, no. FC-121-1030) in the presence of 0.01% digitonin and 0.1% Tween-20 at 37 degree for 30 minutes. The reaction was then cleaned up by DNA Clean and Concentrator-5 Kit (Zymo, no. D4014) and PCR amplified with NEBNext High-Fidelity 2X PCR Master Mix (NEB, no. M0541L) and Nextera Index kit (Illumina, no. 15055289). The resulting libraries were pooled and sequenced on Illumina NextSeq500, according to validated standard operating procedures established by the University of Virginia Genome And Technology Core, RRID: SCR_018883. Oligo sequences refer to **Table S2**.

### Chromatin Endogenous Cleavage (ChEC) assay

#### Cloning

The flexible linker with 6xGly and 3xFLAG was created by gene synthesis (IDT) and cloned into the NheI/BamHI sites of pCW-TetOn-wtCas9 to create pCW3. pCW3-nMN was constructed by insertion of a PCR amplicon encoding the codon optimized deltaSP-nuclease (Addgene #70231) into the HpaI site of pCW3. The ChEC-tagging vector pCW3-nMN-hsORCs was constructed by insertion of a PCR amplicon encoding human ORCs into the AfeI site of pCW3-nMN. Oligo sequences refer to **Table S2**.

#### Stable cell lines

ChEC cell lines expressing MNase-tagged ORC subunits were made with lentivirus by co-transfection of psPAX2, pMD2.G and pCW-nMN-ORC1 (or ORC2 or ORC5) plasmids in the background of ORC1KO-B14, ORC2KOB2 or ORC5KO-24 cells and selected by puromycin. Monoclonal cell population was obtained by limiting dilution.

#### ChEC-seq library construction

For each ChEC experiment, cells were grown in five 10-cm plates. Cells were pelleted at 600 × g at 4□°C for 3□min and washed three times with Washing Buffer (20 mM HEPES-NaOH pH 7.5, 150 mM NaCl, 0.1□mM EGTA, 0.5□mM spermidine, 0.1% BSA, 1 × Roche cOmplete EDTA-free mini protease inhibitors, 1□mM PMSF), centrifuging as above between washes. Cells were resuspended in NE Buffer (20 mM HEPES-KOH pH 7.9, 10 mM KCl, 0.1 mM EGTA, 0.5 mM Spermidine, 0.1% Triton X-100, 20% Glycerol, 1 × Roche cOmplete EDTA-free mini protease inhibitors) and rotated at 4□°C for 10□min. After centrifugation, cells were washed once each with Washing Buffer and Wash Buffer without protease inhibitor and resuspended in Wash Buffer without protease inhibitor. CaCl2 was added to 2□mM and ChEC digests were performed at 25□°C. At each time point, a 150-μl aliquot of the digest was transferred to a tube containing 150□μl 2 × stop buffer (200□mM NaCl, 20□mM EDTA, 4□mM EGTA, 50 µg/mL RNase A, 20 µg /mL Glycogen) and incubated at 37□°C for 30□min. After adding 0.1% SDS, protein was digested with 100□μg proteinase K at 55□°C for 30□min. DNA was extracted one each with an equal volume of phenol and phenol/chloroform/isoamyl alcohol. DNA in aqueous phase was size-selected with 0.45x vol SPRIselect beads and then precipitated with 200 mM NaCl and 2.5 volumes 100% ethanol. Pellets were washed once with 1_ml 70% ethanol, dried, and resuspended in 30_μl 0.1 × TE buffer, pH 8.0. Sequencing libraries was prepared by NEBNext Ultra II DNA Library Prep Kit for Illumina (E7645S).

### RNA-seq library preparation

RNA-seq libraries were prepared as previously(Su et al., 2022). Total RNAs (1_μg) were poly-A selected by NEBNext poly(A) mRNA Magnetic Isolation Module (NEB #E7490) and followed by NEBNext UltraII Directional RNA Kit (NEB #7765). The resulting libraries were pooled and sequenced as paired-end reads on Illumina HiSeq2000 (Novogene) or NextSeq (UVA GATC core facility, RRID: SCR_018883) with >20 million reads per sample.

### HiC-seq library preparation

HiC-seq for cell lines was performed based on HiC2.0 protocol (Belaghzal et al., 2017) with slight modifications. Briefly, cells were crosslinked with 1% formaldehyde for 10 minutes and quenched with 140 mM glycine. Five million crosslinked cells were lysed with 10 mM Tris-HCl, pH 8.0, 10 mM NaCl, 0.2% NP-40. Pelleted nuclei were lysed in 0.1% SDS and quenched by 1% Triton X-100, followed by digestion with DpnII (NEB, no. R0543) at 37 °C overnight in ThermoMixer with constant mixing at 600 RPM. Digested DNA ends were then labeled with biotin-14-dATP (Invitrogen, no. 19524016) by DNA polymerase I, large (Klenow) fragment (NEB, no. M0210) at 37 °C, 1 hour. Labeled DNA was ligated by T4 DNA ligase (Invitrogen, no. 15224017) and incubated with Proteinase K (Invitrogen, no. 25530015) to reverse crosslinking. Resulting DNA was purified and sonicated by Covaris to produce DNA fragments ∼100-400bp. The biotin-labeled DNA fragments were then enriched by MyOne Streptavidin C1 beads (Invitrogen, no. 65001). DNA fragments while on beads were end repaired and made with NEBNext UltraII DNA library prep kit (NEB, no. E7645) with NEBNext Adaptor and Index (PCR amplified 9 cycles). Lastly the libraries were size selected by AMpure XP beads and sequenced on Illumina NovaSeq S4 lane at Novogene. Quality check at various steps was performed according to the HiC2.0 protocol.

### CTCF cut&run library preparation

CUT&RUN Assay Kit (cell signaling technology, no. 86652) was used to perform cut&run according to the manufacturer’s instructions. Anti-CTCF (cell signaling technology, no. 3418) was used.

### ChIP-seq library preparation

ChIP was performed using SimpleChIP® Enzymatic Chromatin IP Kit (Magnetic Beads) (cell signaling technology, no. 9003) according to the manufacturer’s instruction. Antibodies used were Anti-H3K9ME3 (cell signaling technology, no. 13969) and Anti-H3K27ME3 (cell signaling technology, no. 9733). Sequencing libraries was prepared using NEBNext Ultra II DNA Library Prep Kit for Illumina (NEB, no. E7645S) according to manufacturer’s instructions.

### Chromatin loading and western blot

Chromatin fractionation was performed as previously described (PMID 11046155). Antibodies used include α-ORC1 (cell signaling technology, no. 4731), α-ORC2 (Santa Cruz Biotechnology, no. sc-32734), α-ORC3 (Santa Cruz Biotechnology, no. sc-23888), α-ORC4, α-ORC5, α-ORC6 (PMID15944161), α-HSP90 (Santa Cruz Biotechnology, no. sc-13119), α-MCM7 (Santa Cruz Biotechnology, no. sc-9966), α-ORCA (PMID 20932478, generous gift from Supriya G. Prasanth, University of Illinois), α-HP1α (cell signaling technology, no. 2616), α-H3K9ME1 (cell signaling technology, no. 14186), α-H3K9ME2 (cell signaling technology, no. 4658), α-H3K9ME3 (cell signaling technology, no. 13969), α-HBO1 (cell signaling technology, no. 58418), α-H3K27ME3 (cell signaling technology, no. 9733), α-Histone H3 (cell signaling technology, no. 4499), α-NFIA (Apcepta, no. AP20998c), α-CHD7 (cell signaling technology, no. 6505).

### Immunofluorescence

HCT116 cells were incubated in CSK buffer (10 mM Pipes, pH 7.0, 100 mM NaCl, 300 mM sucrose, and 3 mM MgCl_2_) containing 0.5 % Triton X-100 for 5 min on ice before fixation in 4 % Paraformaldehyde in PBS for 10 min. Anti-HP1alpha (1:100, cell signaling technology, no. 2616) was then applied. Image acquisition was performed with Zeiss AxioObserver Z1, 63 X objective.

## Data analysis

### ATAC-seq data analysis

Raw sequence data in FASTQ format were processed as follows: FastQC (v0.11.5) (Andrews, 2010) was used for quality control and sequence data were then mapped to human genome (hg38) using bowtie2 (v2.2.9) (Langdon, 2015). Sam files were converted into bam files using samtools (v1.12) (Li et al., 2009) and only high-quality reads (q-score ≥30) and uniquely mapped reads were retained for subsequent analyses. Bam files were then converted to bed files using samtools. Reads mapped to mitochondrial DNA (chrM) and unplaced or unlocalized contigs were excluded from downstream analyses. Only fragments ≤ 150 bp were retained for ATAC-seq peak calling. Peak calling for ATAC-seq data was performed with MACS2 callpeak function (callpeak -g hs -B --SPMR -q 0.05 --keep-dup 1 -f BEDPE) (v2.1.4) (Zhang et al., 2008).

### RNA-seq data analysis

Raw sequence data in FASTQ format were processed as follows: FastQC (v0.11.5) (Andrews, 2010) was used for quality control and sequence data were then mapped to human genome (hg38) using hisat2 version 2.2.1 (Kim et al., 2019) (hisat2 -t -p 20 −5 20 −3 80). Sam files were converted into bam files using samtools (v1.12) (Li et al., 2009) and only high-quality reads (q-score ≥30) were retained for subsequent analyses. Reads mapped to mitochondrial DNA (chrM) and unplaced or unlocalized contigs were excluded from downstream analyses. HTSeq version 2.0.5 (Putri et al., 2022) was used to calculate the read count for every gene using command *htseq-count -f bam -r name -s no*. The reference genome is hg38 ncbiRefSeq. Differential gene analysis was performed using DESeq2 version 1.42.0 (Love et al., 2014).

### ChEC-seq data analysis

Raw sequence data in FASTQ format were processed as follows: FastQC (v0.11.5) (Andrews, 2010) was used for quality control and sequence data were then mapped to human genome (hg38) using bowtie2 (v2.2.9) (Langdon, 2015). Sam files were converted into bam files using samtools (v1.12) (Li et al., 2009) and only high-quality reads (q-score ≥30) were retained for subsequent analyses. Reads mapped to mitochondrial DNA (chrM) and unplaced or unlocalized contigs were excluded from downstream analyses. Peak calling for ChEC -seq data was performed with MACS2 callpeak function (callpeak -g hs -B --SPMR -q 0.01 --keep-dup 1 -f BAMPE) (v2.1.4) (Zhang et al., 2008), the time 0 sample was used as a background control to account for non-specific cleavage.

### CUT&RUN data analysis

Raw sequence data in FASTQ format were processed using the same pipeline for ChEC-seq data. Peak calling for CTCF was performed with MACS2 (callpeak -g hs -B --SPMR -q 0.05 -- keep-dup 1 -f BAMPE).

### ChIP-seq data analysis

Raw sequence data in FASTQ format were processed using the same pipeline for ChEC-seq data. Peak calling for H3K9me3 and H3K27me3 ChIP-seq data was performed with SICER2 using default parameters (Zang et al., 2009).

### Differential peak analysis

Peaks for different conditions were then merged into a union bed files using Bedtools merge (v2.29.2) (Quinlan and Hall, 2010). Reads count on each of the union peaks were calculated using bedtools intersection function (intersectBed -c) (v2.29.2) (Quinlan and Hall, 2010). Differential peaks were called using DESeq2 version 1.42.0 (Love et al., 2014).

### BART, Permutation test, Motif enrichment analysis

To identify transcription regulator enriched with each list of genomic regions, BART (Binding Analysis for Regulation of Transcription) analysis was performed against publicly available ChIP-seq datasets with BART2 (Wang et al., 2018). To test if two list of genomic regions are overlapped more than random expectations, permutation test is performed with regioneR package with indicated window size (Gel et al., 2016). To identify enriched motifs for CTCF binding sites, Simple Enrichment Analysis in MEME suite was used against JASPAR2022 CORE vertebrates non-redundant v2 motif database.

### HiC-seq data processing and analysis

The paired-end FASTQ data was trimmed to 36bp from 5’ end and then align to the hg19 reference genome using Bowtie for R1 and R2 separately. Then the two mapped SAM files were merged, and the PCR duplication was removed. The reads were then processed following the *HiCorr* pipeline (Lu et al., 2020). All the reads were assigned to the DPNII-digested fragments and reads that within the same fragment was removed. Next, all reads were characterized as ‘inward’, ‘outward’, or ‘same-strand’ based on the strand of R1 and R2. All ‘inward’ reads that had <1kb distance or ‘outward’ reads that had <5kb distance are removed. The remaining ‘inward’ and ‘outward’ reads, together with the ‘same-strand’ reads were merged together as the cis reads. The interchromosomal reads were considered as trans reads. The fragments were then assigned to 5kb anchors to do the following bias correction steps. Next, the data was enhanced with *DeepLoop* universal model and is ready for the downstream analysis (Zhang et al., 2022). All the data were lifted over to hg38 assembly to incorporate with other data analysis. The specific loops called were using the criteria that the |Fold Change| > 2 for the loop strength after *DeepLoop* enhancement in the top 300k loops. PC1 was picked for the principal component analysis of Hi-C data at 250kb resolution.

## Notes

### Competing Interest Statement

The authors have declared no competing interest.

### Summary of Updates

Supplementary Figures have been revised to include new analysis, corresponding texts and methods section are updated.

