## Supplementary Figure for "Regulation of epigenetics and chromosome structure by human ORC2"

**Figure S1. ORC2-dependent changes in H3K9me3 and H3K27me3 distribution. Related to Figure 1.**
(A) Scheme of experiments. (B) Rescue of ORC2 expression in clone 5 comparable to that of WT cells. 1:2 dilution of WT lysate loaded to ensure linearity of ORC2 signal. (C) ORC1, ORC4 and ORC5 on the chromatin disappear when ORC2 is deleted, and reappear in the rescue cell line. (D) H3K27me3 ChIP-seq peaks altered in knockout cells and rescued by reintroduction of ORC2. (E) Example H3K9me3 and H3K27me3 peaks (n = 2). The appearance of H3K9me3 is accompanied by a decrease of H3K27me3 in the ORC2 knockout. (F-G) Anti-correlation of H3K9me3 signal and H3K27me3 signal when ORC2 is deleted.

**Figure S2. ORC2-dependent chromatin accessibility changes suggest its role in chromatin repression. Related to Figure 2.**
(A) Sequence length distribution of ATAC-seq DNA shows DNA fragments that are nucleosomal and subnucleosomal. (B-C) Heatmap of (B) H3K9me3 and (C) H3K27me3 ChIP-seq signal (n = 2) in three cell lines centered on ATAC-up peaks. (D-E) Heatmap of (D) H3K9me3 and (E) H3K27me3 ChIP-seq signal (n = 2) in three cell lines centered on ATAC-down peaks.

**Figure S3. ORC2-dependent chromatin accessibility changes are accompanied by changes in gene expression. Related to Figure 3.**
(A) Gene Set Enrichment analysis of genes dysregulated in the ORC2KO relative to WT. Red: Top 10 Gene Ontologies (Molecular Function) enriched in up-regulated genes. Blue: Top 10 Gene Ontologies (Molecular Function) enriched in down-regulated genes. (B) Same as (A) except for genes that are altered in the ORC2 rescued cells relative to the knockout.

**Figure S4. ORC2-dependent chromatin accessibility and gene expression changes are specific to ORC2 and not ORC1 or ORC5. Related to Figure 4.**
(A) Expression changes of genes in ORC1 KO cells correlated with change of ATAC accessibility in the TSS region. (B) Example ATAC sites increased only by loss of ORC2, but not ORC1 or ORC5: chr2: 28,116,699-28,117,951; chr17: 2,067,878-2,068,738; chr1: 26,772,241-26,773,528; chr16: 72,924,642-72,925,575. (C) Example ATAC sites decreased only by loss of ORC1, but not ORC2 or ORC5: chr5: 65,563,033-65,563,384; chr1: 193,121,347-193,122,746; chr1: 193,058,245-193,061,044; chr22 :19,963,039-19,963,340. (D) Example ATAC sites decrease by loss of ORC5, but not ORC1 or ORC5 (left two loci), or also decreased by loss of ORC1 (third column) or increased by loss of ORC2: chr2: 40,434,830-40,435,299; chr17: 68,383,773-68,384,419; chr7: 46,764,069-46,764,687; chr5: 33,049,244-33,049,948. (E) Gene Set Enrichment analysis of genes dysregulated in the ORC1KO relative to WT. (F) Same as E) except for gene dysregulated in ORC5 KO relative to WT.

**Figure S5. Binding sites of ORC1, ORC2 and ORC5. Related to Figure 5.**
(A-C) Western blots show expression of the micrococcal nuclease tagged ORC subunit has comparable expression to that of the endogenous subunit. (D) Western blots of subcellular fractions. Restoration of MN-ORC2 in the ORC2D cells restores the chromatin binding of ORC1, ORC3-5 and MCM7, showing that the MN-ORC2 is functional. (E) Venn diagram to identify the reproducible binding sites of indicated subunits from two independent clones. Only the reproducible sites are used for downstream analysis. (F) Circos plot shows binding density of the three ORC subunits to the genome. (G) Example sites bound by all three or subsets of ORC subunits. The ATAC-seq plots show the different changes in chromatin accessibility at the same sites when ORC2 is deleted. Red box: ATAC accessibility increases in ORC2KO at sites bound by all three subunits (Chr 1) or only ORC2 (Chr 15). (H) Enrichment relative to random expectation (expressed as Z score of 1000 random permutations) of ORC2 ChEC-seq sites with indicated ATAC-sites and histone modifications (ENCODE HCT116). (I-J) Venn diagrams to show overlap

of ORC2 binding sites with ORC2-dependent ATAC-up or ATAC-down sites. (K) ORC2 bound sites show a regional decrease of H3K9me3 when ORC2 is deleted. (L-M) BART analysis of ORC1-bound and ORC5-bound sites to identify the locally co-enriched transcription factors. (N) Overlap between ORC2-regulated genes identified from RNA-seq (Fig. 4) and ORC2 ChEC-seq sites. (O) ORC2 ChEC-seq signals over ORC2-regulated genes identified from RNA-seq. (P) Similar to Fig. 5B, but using union peaks from each ORC1/ORC2/ORC5 ChEC-seq datasets.

**Figure S6. ORC2 is required for specification of some chromosome compartments and some chromatin loops. Related to Figure 6.**
(A) Table showing total number of reads in Hi-C experiments and the number of cis- and transinteractions discovered. (B) Scatter plot of compartment changes (defined by PC1 value) in the KO vs WT or KO vs rescue. (C) Enrichment of ORC2 binding sites across the region centered on compartments that are changed in KO relative to the WT by permutation test (n = 1000). Red: compartments that are decompacted (B to A) when ORC2 is deleted. Blue: compartments that change in the reverse direction (A to B). (D) Compartment designations (Red: A; Blue: B) of two chromosomes in the three cell lines. (E) Scatter plot of loop strength in the KO vs WT or KO vs rescue. The red and blue dots indicate loops that were significantly changed (cut-off: fold change > 2). (F) Enrichment of Loop-up sites (in the KO relative to the WT) with indicated ATAC or H3K9me3 changes in the KO relative to the WT by permutation test (n = 1000).

**Figure S7. A functional interaction between ORC2 and CTCF driven by local accessibility. Related to Figure 7.**
(A) Overlap of all CTCF binding sites with ORC2 binding sites. (B) More CTCF-up sites (increased in the ORC2 KO) overlap with ORC2 binding sites. (C) Same as Figure 7(E) except for the NFIA locus.

**Figure S1.**

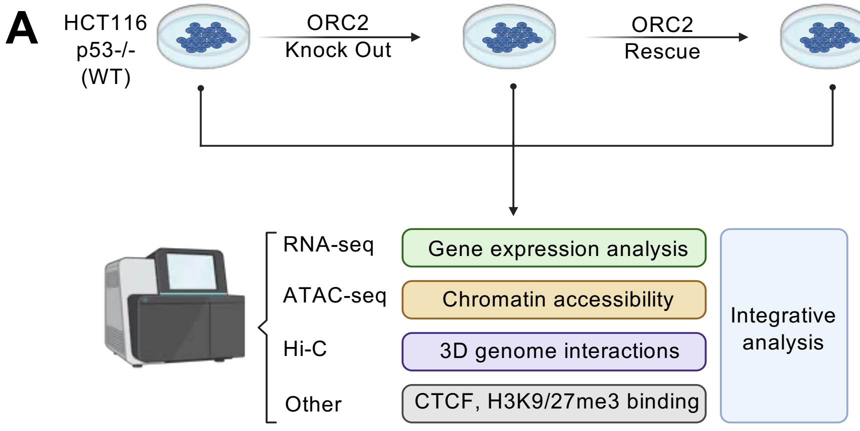


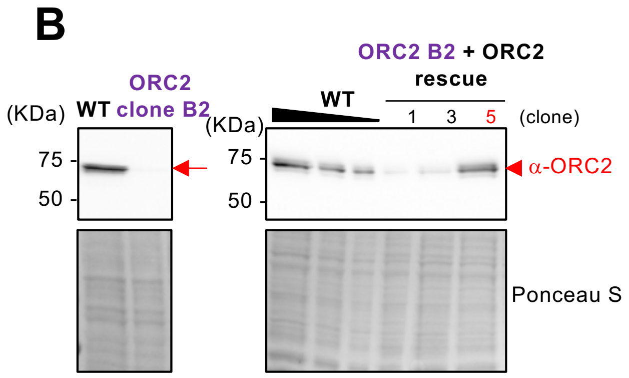


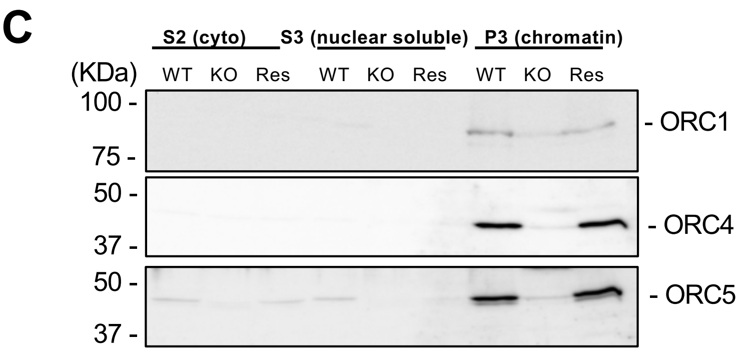


**D** *H3K27me3 ChIP-seq*

| # of peaks | Up in ORC2 KO | Down in ORC2 KO |
|---|---|---|
| ORC2 KO vs WT | 3474 | 5362 |
| Rescued | 984 (28%) | 623 (12%) |

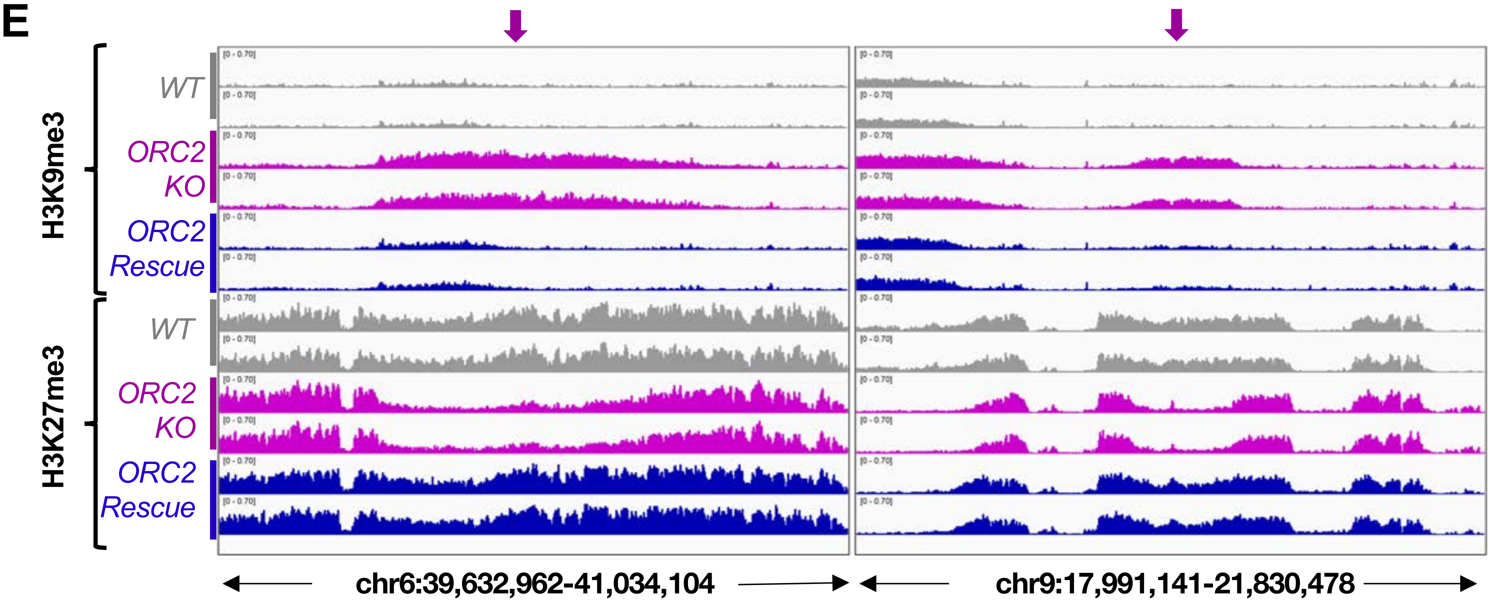


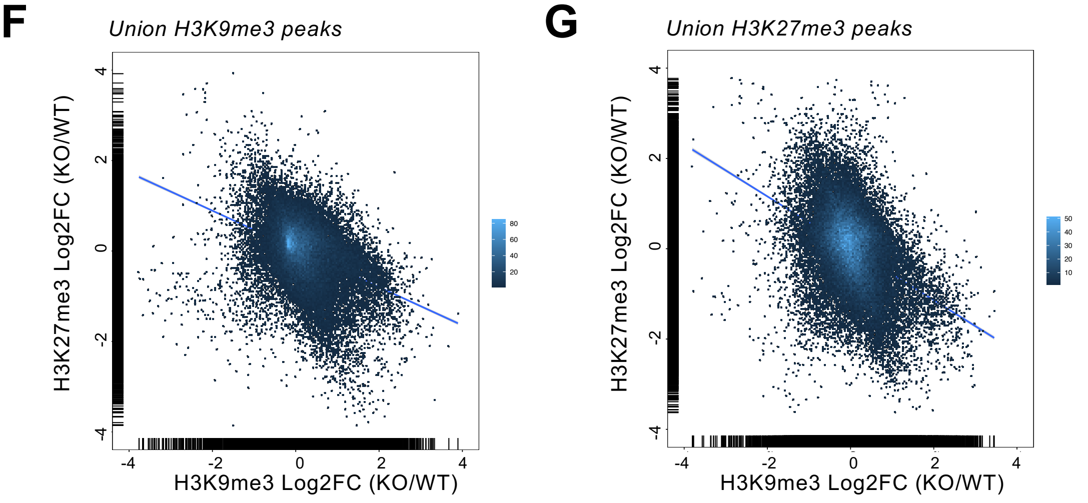

Figure S2.

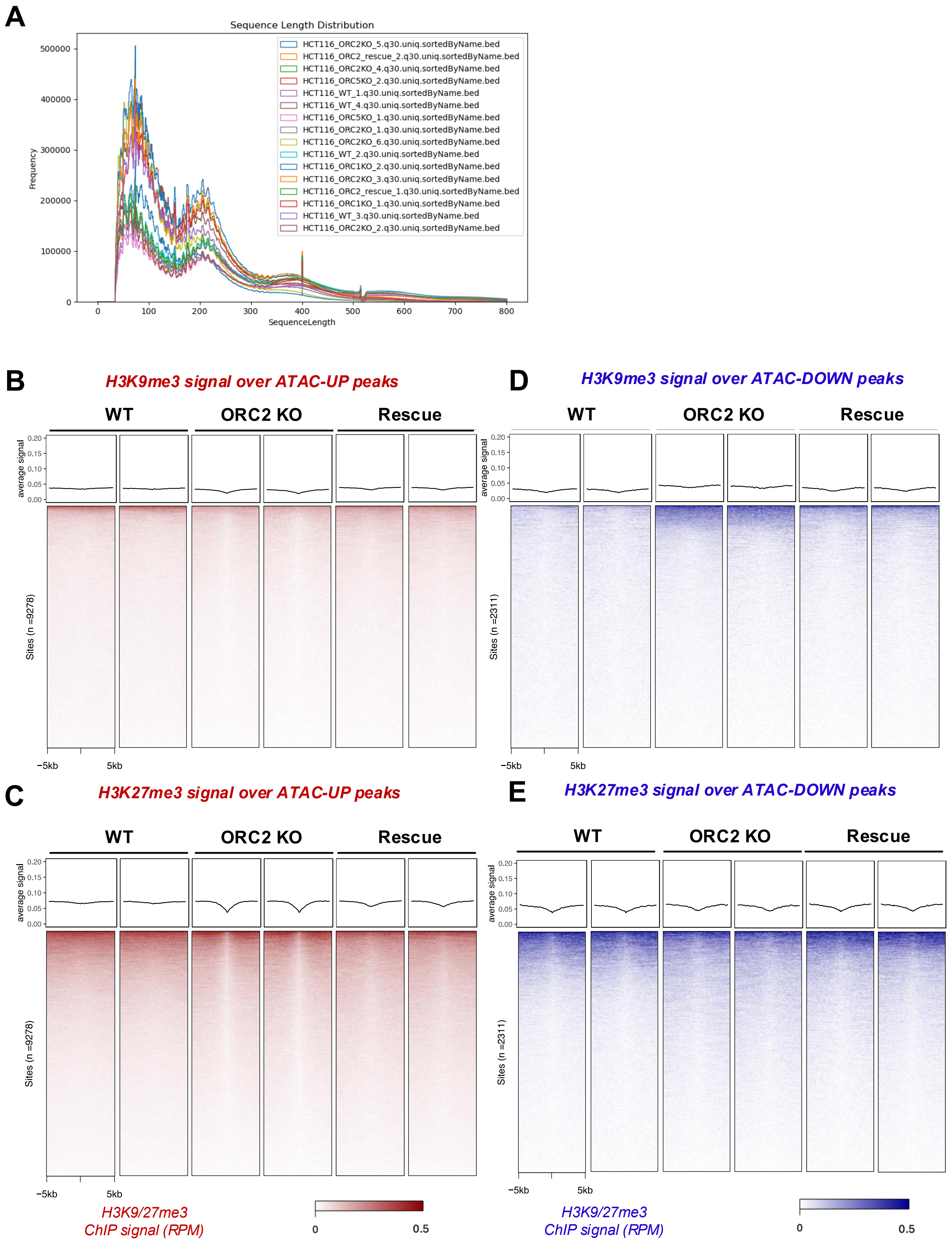
A
Sequence Length Distribution
HCT116_ORC2KO_5.q30.uniq.sortedByName.bed
HCT116_ORC2_rescue_2.q30.uniq.sortedByName.bed
HCT116_ORC2KO_4.q30.uniq.sortedByName.bed
HCT116_ORC5KO_2.q30.uniq.sortedByName.bed
HCT116_WT_1.q30.uniq.sortedByName.bed
HCT116_WT_4.q30.uniq.sortedByName.bed
HCT116_ORC5KO_1.q30.uniq.sortedByName.bed
HCT116_ORC2KO_1.q30.uniq.sortedByName.bed
HCT116_ORC2KO_6.q30.uniq.sortedByName.bed
HCT116_WT_2.q30.uniq.sortedByName.bed
HCT116_ORC1KO_2.q30.uniq.sortedByName.bed
HCT116_ORC2KO_3.q30.uniq.sortedByName.bed
HCT116_ORC2_rescue_1.q30.uniq.sortedByName.bed
HCT116_ORC1KO_1.q30.uniq.sortedByName.bed
HCT116_WT_3.q30.uniq.sortedByName.bed
HCT116_ORC2KO_2.q30.uniq.sortedByName.bed
Frequency
SequenceLength
B
H3K9me3 signal over ATAC-UP peaks
D
H3K9me3 signal over ATAC-DOWN peaks
WT
ORC2 KO
Rescue
average signal
Sites (n =9278)
Sites (n =2311)
−5kb
5kb
C
H3K27me3 signal over ATAC-UP peaks
E
H3K27me3 signal over ATAC-DOWN peaks
H3K9/27me3
ChIP signal (RPM)
0
0.5

**Figure S3.**

**A** ORC2 KO vs WT

| Pathway | NES | pval | padj |
|---|---|---|---|
| GOMF_STRUCTURAL_CONSTITUENT_OF_SKIN_EPIDERMIS | 2.35 | 4.4e−09 | 3.0e−07 |
| GOMF_COLLAGEN_BINDING | 2.22 | 6.4e−08 | 3.3e−06 |
| GOMF_EXTRACELLULAR_MATRIX_STRUCTURAL_CONSTITUENT | 2.13 | 1.9e−09 | 1.7e−07 |
| GOMF_SODIUM_CHANNEL_ACTIVITY | 2.01 | 9.9e−05 | 2.2e−03 |
| GOMF_INTEGRIN_BINDING | 1.99 | 1.8e−07 | 8.4e−06 |
| GOMF_EXTRACELLULAR_MATRIX_STRUCTURAL_ CONSTITUENT_CONFERRING_TENSILE_STRENGTH | 1.99 | 6.9e−05 | 1.8e−03 |
| GOMF_STRUCTURAL_CONSTITUENT_OF_MUSCLE | 1.99 | 7.5e−05 | 1.8e−03 |
| GOMF_EXTRACELLULAR_MATRIX_BINDING | 1.98 | 9.9e−05 | 2.2e−03 |
| GOMF_AMINOACYLTRANSFERASE_ACTIVITY | 1.95 | 2.3e−04 | 4.3e−03 |
| GOMF_MONOATOMIC_CATION_CHANNEL_ACTIVITY | 1.94 | 2.0e−09 | 1.7e−07 |
| GOMF_HISTONE_BINDING | −2.18 | 3.3e−09 | 2.5e−07 |
| GOMF_ATP_DEPENDENT_ACTIVITY_ACTING_ON_RNA | −2.19 | 2.9e−05 | 8.9e−04 |
| GOMF_NUCLEOSOME_BINDING | −2.20 | 3.6e−05 | 1.0e−03 |
| GOMF_MIRNA_BINDING | −2.21 | 3.0e−04 | 5.0e−03 |
| GOMF_CATALYTIC_ACTIVITY_ACTING_ON_DNA | −2.24 | 2.0e−11 | 4.0e−09 |
| GOMF_REGULATORY_RNA_BINDING | −2.25 | 6.2e−05 | 1.6e−03 |
| GOMF_DNA_SECONDARY_STRUCTURE_BINDING | −2.26 | 1.2e−04 | 2.5e−03 |
| GOMF_ATP_DEPENDENT_ACTIVITY_ACTING_ON_DNA | −2.31 | 1.3e−08 | 7.5e−07 |
| GOMF_HELICASE_ACTIVITY | −2.39 | 8.9e−11 | 1.1e−08 |
| GOMF_SINGLE_STRANDED_DNA_BINDING | −2.52 | 1.6e−11 | 4.0e−09 |

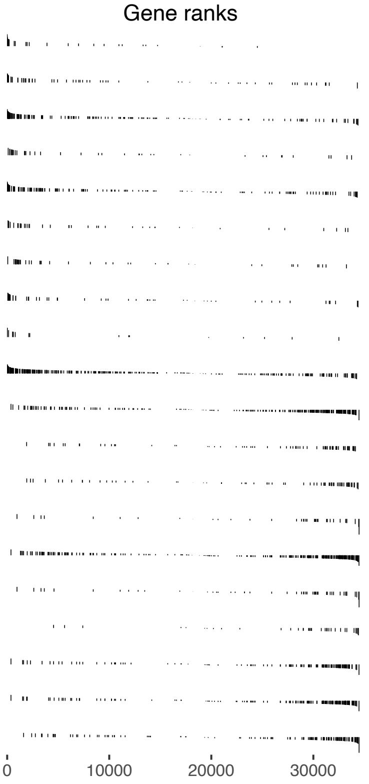


**B** Rescue vs ORC2 KO

| Pathway | NES | pval | padj |
|---|---|---|---|
| GOMF_HELICASE_ACTIVITY | 2.22 | $8.1\cdot10^{-9}$ | $6.8\cdot10^{-7}$ |
| GOMF_ATP_DEPENDENT_ACTIVITY_ACTING_ON_DNA | 2.11 | $5.8\cdot10^{-7}$ | $2.6\cdot10^{-5}$ |
| GOMF_CATALYTIC_ACTIVITY_ACTING_ON_DNA | 2.08 | $7.0\cdot10^{-9}$ | $6.7\cdot10^{-7}$ |
| GOMF_HISTONE_BINDING | 2.08 | $2.0\cdot10^{-9}$ | $2.2\cdot10^{-7}$ |
| GOMF_CATALYTIC_ACTIVITY_ACTING_ON_RNA | 2.07 | $2.7\cdot10^{-13}$ | $1.0\cdot10^{-10}$ |
| GOMF_STRUCTURAL_CONSTITUENT_OF_NUCLEAR_PORE | 2.02 | $4.6\cdot10^{-4}$ | $6.1\cdot10^{-3}$ |
| GOMF_ATP_DEPENDENT_ACTIVITY_ACTING_ON_RNA | 2.00 | $3.5\cdot10^{-5}$ | $7.3\cdot10^{-4}$ |
| GOMF_DNA_REPLICATION_ORIGIN_BINDING | 2.00 | $5.3\cdot10^{-4}$ | $6.7\cdot10^{-3}$ |
| GOMF_PROMOTER_SPECIFIC_CHROMATIN_BINDING | 2.00 | $8.6\cdot10^{-5}$ | $1.5\cdot10^{-3}$ |
| GOMF_FOUR_WAY_JUNCTION_DNA_BINDING | 1.99 | $6.6\cdot10^{-4}$ | $8.0\cdot10^{-3}$ |
| GOMF_INTEGRIN_BINDING | −2.03 | $1.4\cdot10^{-7}$ | $7.5\cdot10^{-6}$ |
| GOMF_ACTIN_BINDING | −2.04 | $1.8\cdot10^{-14}$ | $1.4\cdot10^{-11}$ |
| GOMF_STRUCTURAL_CONSTITUENT_OF_MUSCLE | −2.05 | $5.9\cdot10^{-6}$ | $1.7\cdot10^{-4}$ |
| GOMF_R_SMAD_BINDING | −2.07 | $3.3\cdot10^{-6}$ | $9.8\cdot10^{-5}$ |
| GOMF_FIBRONECTIN_BINDING | −2.09 | $2.8\cdot10^{-6}$ | $8.9\cdot10^{-5}$ |
| GOMF_LAMININ_BINDING | −2.13 | $3.3\cdot10^{-6}$ | $9.8\cdot10^{-5}$ |
| GOMF_EXTRACELLULAR_MATRIX_STRUCTURAL_CONSTITUENT | −2.20 | $8.5\cdot10^{-11}$ | $2.1\cdot10^{-8}$ |
| GOMF_STRUCTURAL_CONSTITUENT_OF_SKIN_EPIDERMIS | −2.23 | $3.9\cdot10^{-8}$ | $2.7\cdot10^{-6}$ |
| GOMF_COLLAGEN_BINDING | −2.30 | $1.2\cdot10^{-9}$ | $1.8\cdot10^{-7}$ |
| GOMF_EXTRACELLULAR_MATRIX_BINDING | −2.31 | $1.5\cdot10^{-9}$ | $1.9\cdot10^{-7}$ |

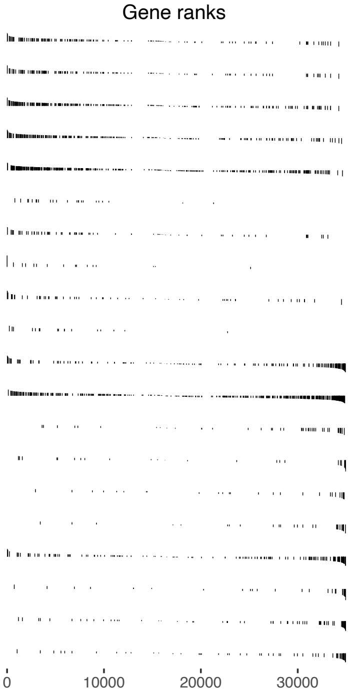

**Figure S4.**

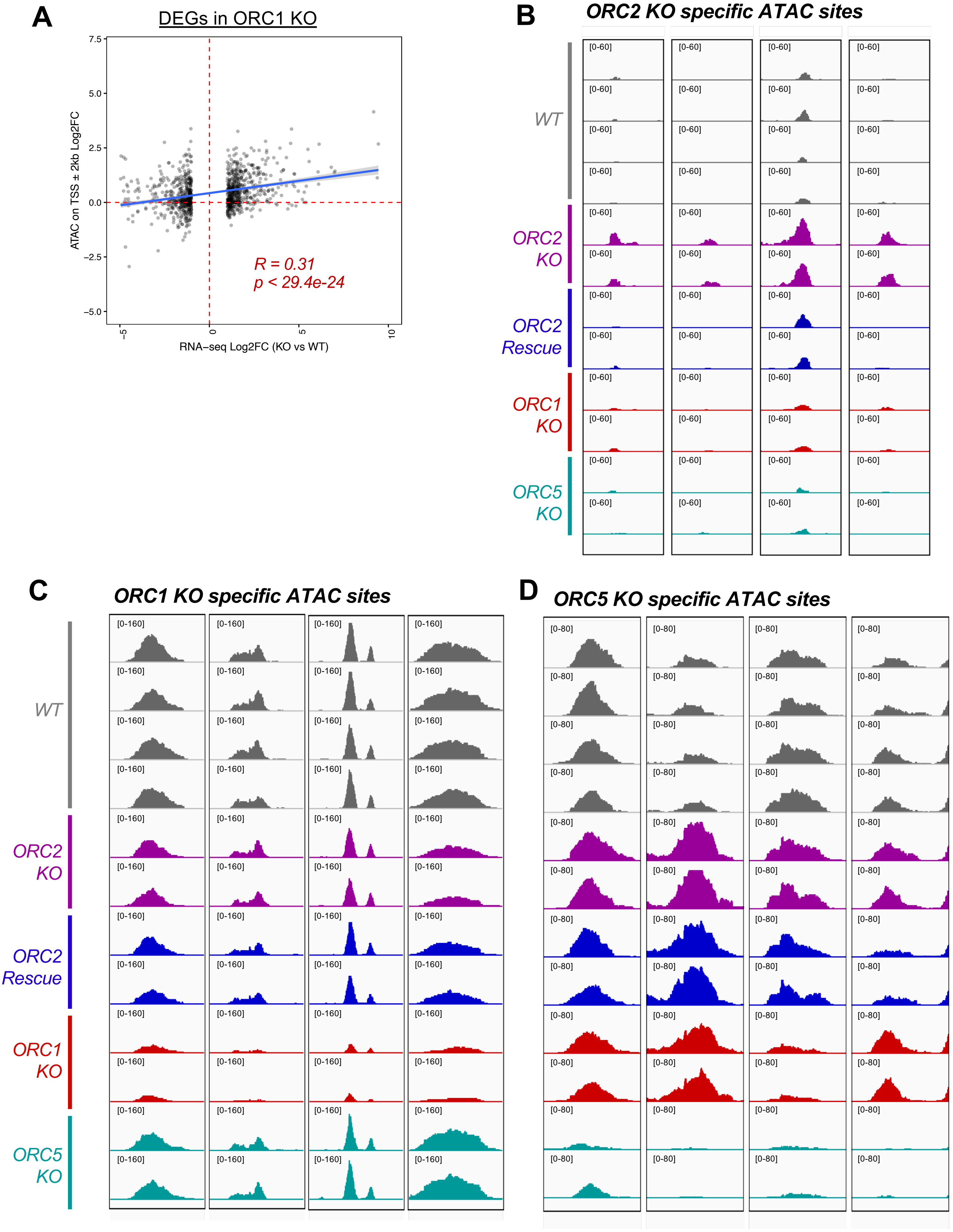
A
DEGs in ORC1 KO
7.5
5.0
2.5
0.0
−2.5
−5.0
ATAC on TSS ± 2kb Log2FC
−5
0
5
10
RNA−seq Log2FC (KO vs WT)
R = 0.31
p < 29.4e-24
B
ORC2 KO specific ATAC sites
[0-60]
WT
ORC2 KO
ORC2 Rescue
ORC1 KO
ORC5 KO
C
ORC1 KO specific ATAC sites
[0-160]
WT
ORC2 KO
ORC2 Rescue
ORC1 KO
ORC5 KO
D
ORC5 KO specific ATAC sites
[0-80]

**Figure S4. (cont.)**

**E**

ORC1 KO vs WT

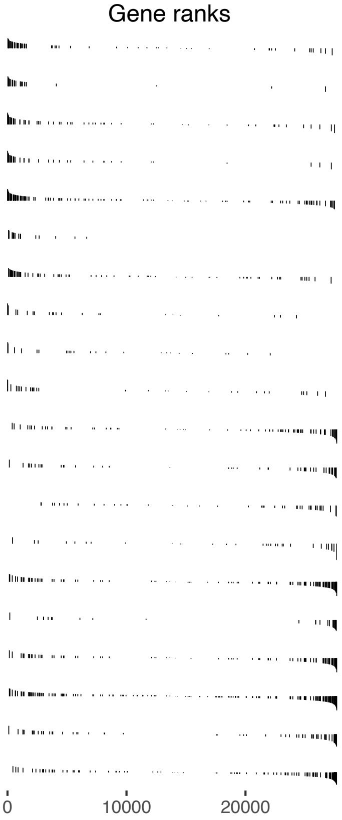


| Pathway | NES | pval | padj |
|---|---|---|---|
| GOMF_DNA_HELICASE_ACTIVITY | 2.14 | $2.9 \cdot 10^{-7}$ | $5.4 \cdot 10^{-5}$ |
| GOMF_SINGLE_STRANDED_DNA_HELICASE_ACTIVITY | 2.09 | $9.1 \cdot 10^{-6}$ | $7.6 \cdot 10^{-4}$ |
| GOMF_PROTEIN_FOLDING_CHAPERONE | 2.01 | $1.4 \cdot 10^{-5}$ | $9.5 \cdot 10^{-4}$ |
| GOMF_ATP_DEPENDENT_PROTEIN_FOLDING_CHAPERONE | 2.01 | $1.4 \cdot 10^{-5}$ | $9.5 \cdot 10^{-4}$ |
| GOMF_UNFOLDED_PROTEIN_BINDING | 2.00 | $8.2 \cdot 10^{-7}$ | $1.2 \cdot 10^{-4}$ |
| GOMF_DNA_REPLICATION_ORIGIN_BINDING | 1.95 | $4.5 \cdot 10^{-5}$ | $2.6 \cdot 10^{-3}$ |
| GOMF_MICROTUBULE_MOTOR_ACTIVITY | 1.93 | $2.9 \cdot 10^{-5}$ | $1.8 \cdot 10^{-3}$ |
| GOMF_MHC_CLASS_II_PROTEIN_COMPLEX_BINDING | 1.92 | $7.3 \cdot 10^{-4}$ | $1.7 \cdot 10^{-2}$ |
| GOMF_STRUCTURAL_CONSTITUENT_OF_SKIN_EPIDERMIS | 1.83 | $3.1 \cdot 10^{-3}$ | $4.7 \cdot 10^{-2}$ |
| GOMF_PROTEIN_CARRIER_CHAPERONE | 1.82 | $1.9 \cdot 10^{-3}$ | $3.1 \cdot 10^{-2}$ |
| GOMF_TRNA_BINDING | −1.78 | $7.8 \cdot 10^{-4}$ | $1.7 \cdot 10^{-2}$ |
| GOMF_TRANSLATION_INITIATION_FACTOR_ACTIVITY | −1.78 | $6.4 \cdot 10^{-4}$ | $1.5 \cdot 10^{-2}$ |
| GOMF_PROTEIN_KINASE_A_BINDING | −1.81 | $4.2 \cdot 10^{-4}$ | $1.4 \cdot 10^{-2}$ |
| GOMF_PROTEIN_TYROSINE_KINASE_ACTIVATOR_ACTIVITY | −1.84 | $2.5 \cdot 10^{-4}$ | $9.2 \cdot 10^{-3}$ |
| GOMF_TRANSLATION_REGULATOR_ACTIVITY_NUCLEIC_ACID_BINDING | −1.87 | $6.9 \cdot 10^{-6}$ | $7.4 \cdot 10^{-4}$ |
| GOMF_MRNA_5_UTR_BINDING | −1.87 | $4.7 \cdot 10^{-4}$ | $1.4 \cdot 10^{-2}$ |
| GOMF_TRANSLATION_FACTOR_ACTIVITY_RNA_BINDING | −1.94 | $9.0 \cdot 10^{-6}$ | $7.6 \cdot 10^{-4}$ |
| GOMF_TRANSLATION_REGULATOR_ACTIVITY | −1.98 | $4.0 \cdot 10^{-9}$ | $1.5 \cdot 10^{-6}$ |
| GOMF_RRNA_BINDING | −2.01 | $3.8 \cdot 10^{-6}$ | $4.7 \cdot 10^{-4}$ |
| GOMF_STRUCTURAL_CONSTITUENT_OF_RIBOSOME | −2.59 | $7.7 \cdot 10^{-33}$ | $5.7 \cdot 10^{-30}$ |

**F**

ORC5 KO vs WT

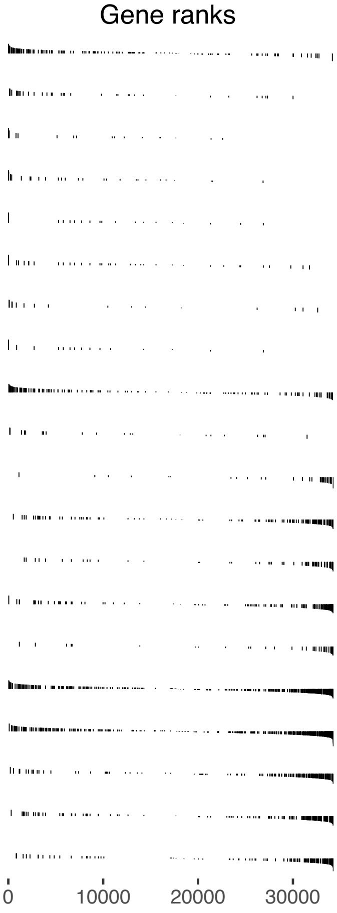


| Pathway | NES | pval | padj |
|---|---|---|---|
| GOMF_EXTRACELLULAR_MATRIX_STRUCTURAL_CONSTITUENT | 2.27 | $2.0 \cdot 10^{-10}$ | $1.0 \cdot 10^{-8}$ |
| GOMF_EXTRACELLULAR_MATRIX_STRUCTURAL_ CONSTITUENT_CONFERRING_TENSILE_STRENGTH | 2.08 | $3.3 \cdot 10^{-5}$ | $4.1 \cdot 10^{-4}$ |
| GOMF_SIALIC_ACID_BINDING | 2.02 | $1.1 \cdot 10^{-4}$ | $1.2 \cdot 10^{-3}$ |
| GOMF_STRUCTURAL_CONSTITUENT_OF_SKIN_EPIDERMIS | 2.01 | $3.1 \cdot 10^{-4}$ | $2.7 \cdot 10^{-3}$ |
| GOMF_RETINAL_BINDING | 1.91 | $6.2 \cdot 10^{-4}$ | $4.7 \cdot 10^{-3}$ |
| GOMF_ISOPRENOID_BINDING | 1.84 | $1.6 \cdot 10^{-3}$ | $1.1 \cdot 10^{-2}$ |
| GOMF_INSULIN_LIKE_GROWTH_FACTOR_BINDING | 1.80 | $4.9 \cdot 10^{-3}$ | $2.5 \cdot 10^{-2}$ |
| GOMF_RETINOL_BINDING | 1.79 | $1.7 \cdot 10^{-3}$ | $1.1 \cdot 10^{-2}$ |
| GOMF_GROWTH_FACTOR_BINDING | 1.77 | $7.0 \cdot 10^{-5}$ | $8.0 \cdot 10^{-4}$ |
| GOMF_NUCLEAR_STEROID_RECEPTOR_ACTIVITY | 1.74 | $6.0 \cdot 10^{-3}$ | $2.8 \cdot 10^{-2}$ |
| GOMF_SNORNA_BINDING | −2.28 | $2.9 \cdot 10^{-8}$ | $8.5 \cdot 10^{-7}$ |
| GOMF_ATP_DEPENDENT_ACTIVITY_ACTING_ON_DNA | −2.32 | $2.2 \cdot 10^{-11}$ | $1.4 \cdot 10^{-9}$ |
| GOMF_DNA_HELICASE_ACTIVITY | −2.34 | $1.9 \cdot 10^{-9}$ | $8.0 \cdot 10^{-8}$ |
| GOMF_UNFOLDED_PROTEIN_BINDING | −2.35 | $1.1 \cdot 10^{-11}$ | $7.4 \cdot 10^{-10}$ |
| GOMF_ATP_DEPENDENT_PROTEIN_FOLDING_CHAPERONE | −2.39 | $3.6 \cdot 10^{-10}$ | $1.7 \cdot 10^{-8}$ |
| GOMF_ATP_HYDROLYSIS_ACTIVITY | −2.39 | $6.6 \cdot 10^{-24}$ | $2.5 \cdot 10^{-21}$ |
| GOMF_MRNA_BINDING | −2.39 | $5.9 \cdot 10^{-24}$ | $2.5 \cdot 10^{-21}$ |
| GOMF_RIBONUCLEOPROTEIN_COMPLEX_BINDING | −2.44 | $1.8 \cdot 10^{-15}$ | $2.0 \cdot 10^{-13}$ |
| GOMF_HELICASE_ACTIVITY | −2.45 | $2.4 \cdot 10^{-16}$ | $3.1 \cdot 10^{-14}$ |
| GOMF_SINGLE_STRANDED_DNA_BINDING | −2.58 | $5.0 \cdot 10^{-17}$ | $7.6 \cdot 10^{-15}$ |

**Figure S5.**

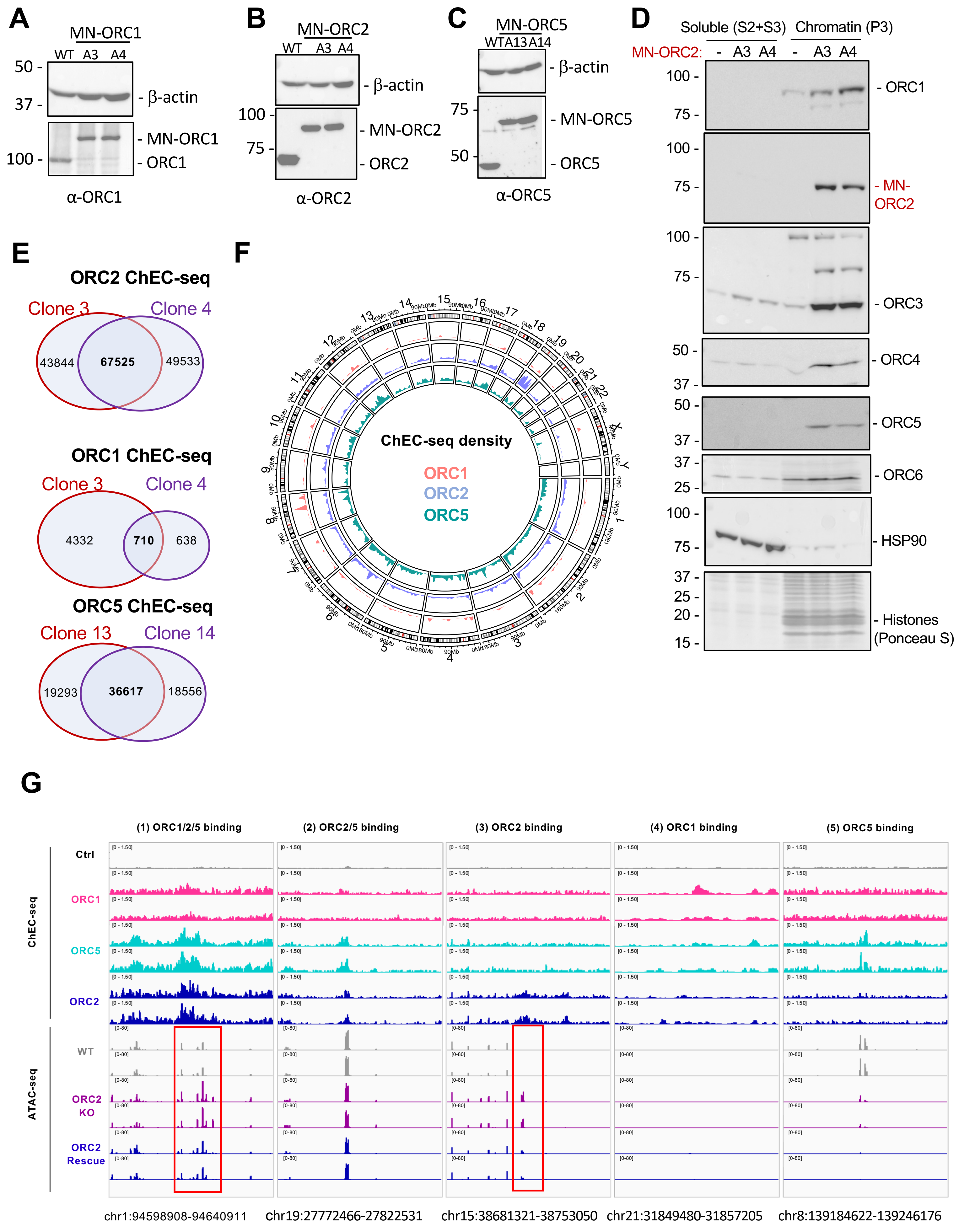
A
MN-ORC1
WT A3 A4
50 -
37 -
100 -
- β-actin
- MN-ORC1
- ORC1
α-ORC1
B
MN-ORC2
WT A3 A4
100 -
75 -
- β-actin
- MN-ORC2
- ORC2
α-ORC2
C
MN-ORC5
WT A13 A14
75 -
50 -
- β-actin
- MN-ORC5
- ORC5
α-ORC5
D
Soluble (S2+S3) Chromatin (P3)
MN-ORC2: - A3 A4 - A3 A4
- ORC1
- MN-ORC2
- ORC3
- ORC4
- ORC5
- ORC6
- HSP90
- Histones (Ponceau S)
E
ORC2 ChEC-seq
Clone 3 Clone 4
43844 67525 49533
ORC1 ChEC-seq
Clone 3 Clone 4
4332 710 638
ORC5 ChEC-seq
Clone 13 Clone 14
19293 36617 18556
F
ChEC-seq density
ORC1
ORC2
ORC5
G
(1) ORC1/2/5 binding
(2) ORC2/5 binding
(3) ORC2 binding
(4) ORC1 binding
(5) ORC5 binding
ChEC-seq
Ctrl
ORC1
ORC5
ORC2
ATAC-seq
WT
ORC2 KO
ORC2 Rescue
chr1:94598908-94640911
chr19:27772466-27822531
chr15:38681321-38753050
chr21:31849480-31857205
chr8:139184622-139246176

Figure S5. (cont.)

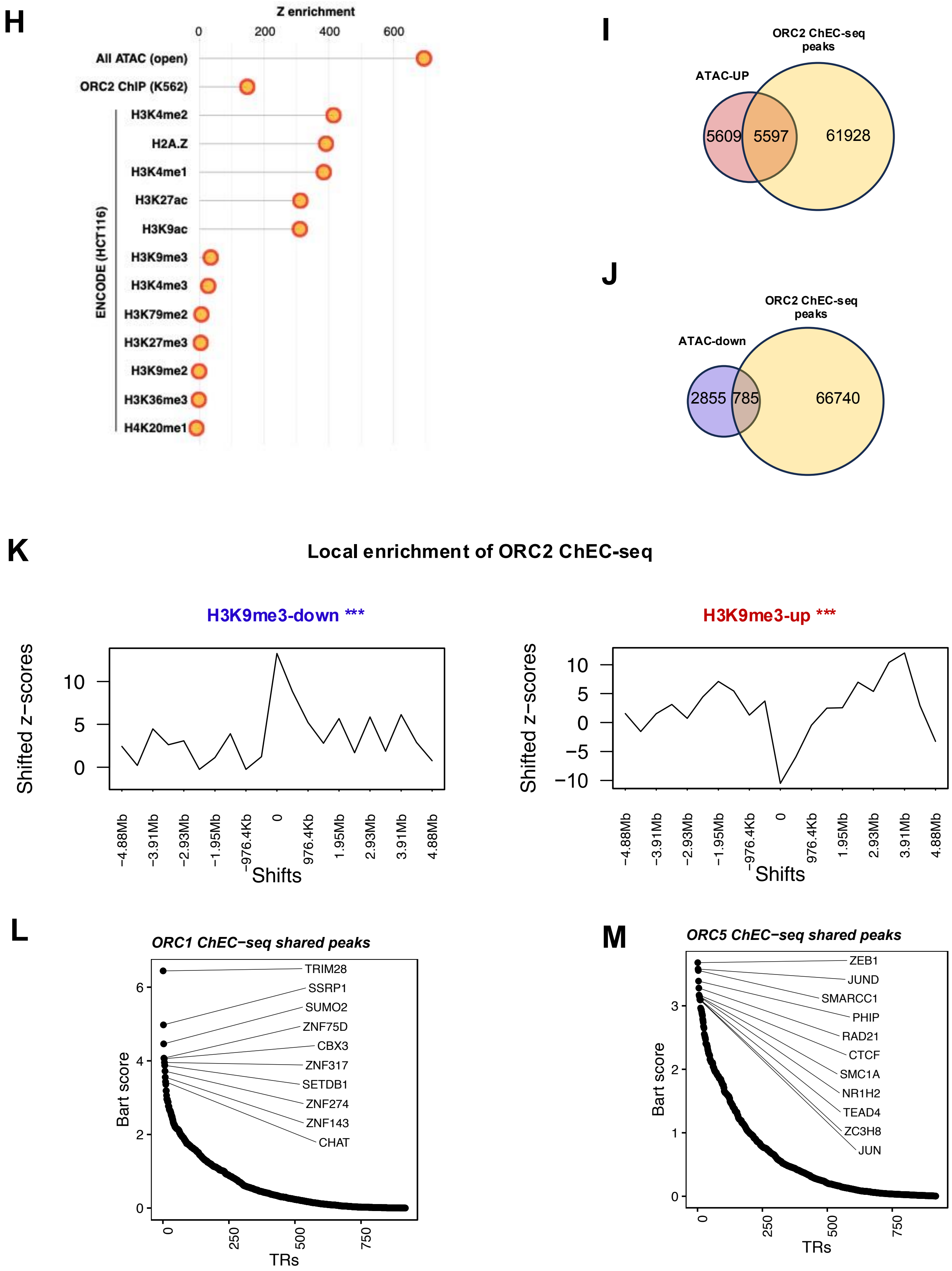

**N** ORC2-regulated genes (mapped to gene body extend 4k) overlapped with ORC2 binding sites

| | RNA-up in ORC2Δ | RNA-down in ORC2Δ | All genes |
|---|---|---|---|
| Number of genes | 824 | 236 | 53598 |
| overlapped with ORC2 | 581 | 137 | 18890 |
| overlapped with ORC2 (%) | 71% | 58% | 35% |

**P** Overlap between ORC1/2/5 binding sites (union peaks)

| # of peaks | ORC2 ChEC | ORC1 ChEC | ORC5 ChEC |
|---|---|---|---|
| ORC2 ChEC | 157897 | - | - |
| ORC1 ChEC | 3765 | 5667 | - |
| ORC5 ChEC | 46429 | 4591 | 73094 |

**O**

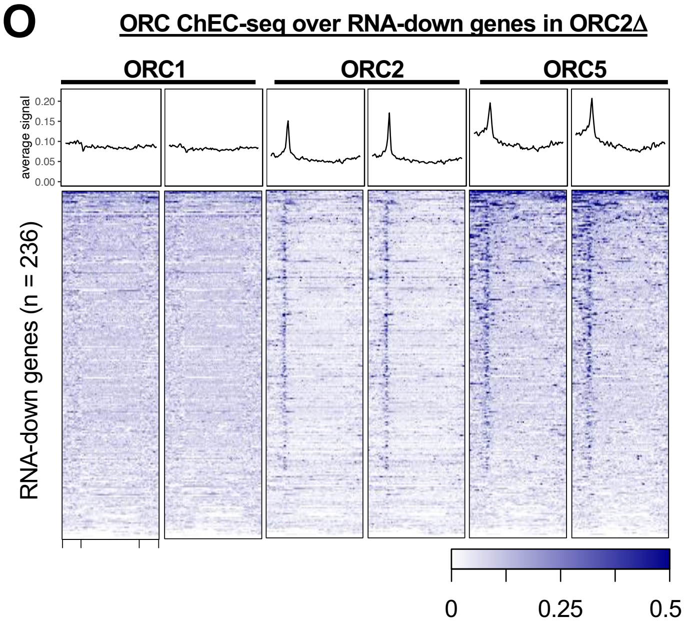


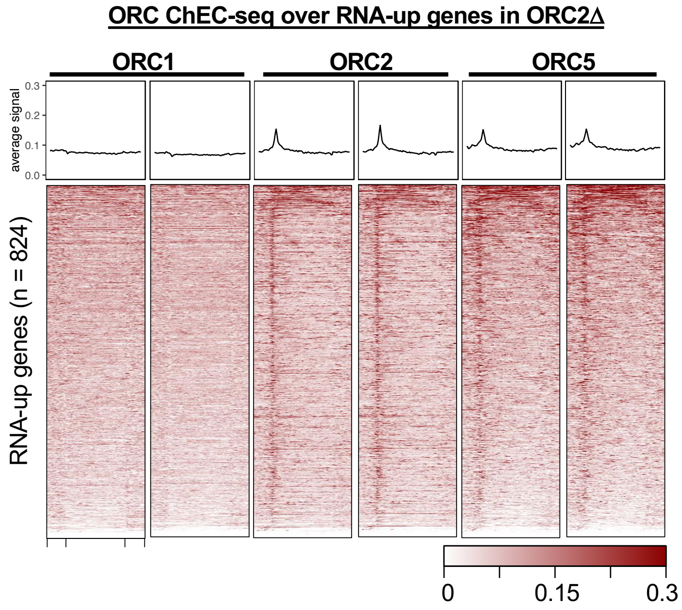

**Figure S6.**

**A**

| | Total # Reads | Cis interactions | Cis interactions within 2M | Trans interactions |
|---|---|---|---|---|
| WT | 880,708,479 | 180,936,973 | 103,431,299 | 108,420,833 |
| ORC2 KO | 888,054,145 | 203,266,322 | 140,804,491 | 77,046,271 |
| Rescue | 767,885,482 | 188,017,391 | 133,410,515 | 71,517,363 |

**B**

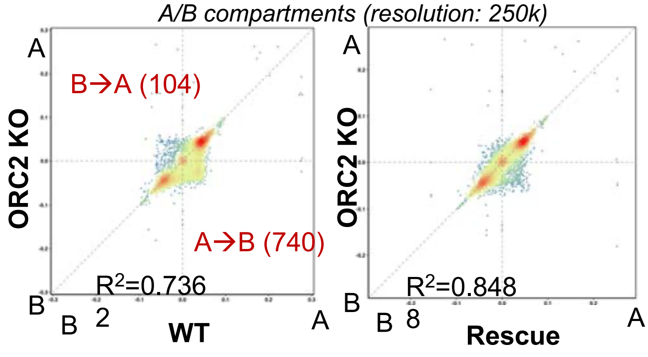


**C**

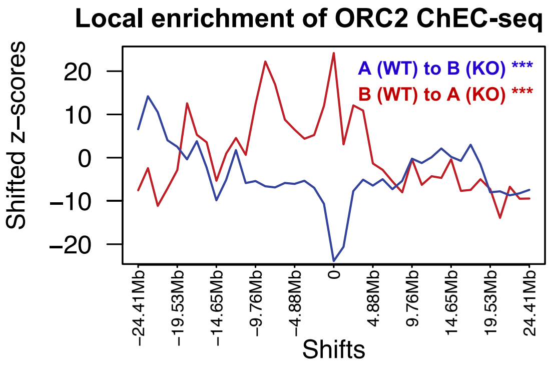


**D**

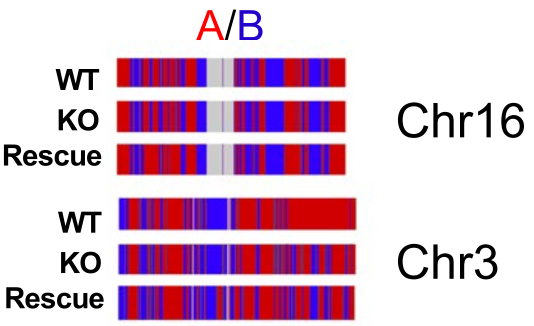


**E**

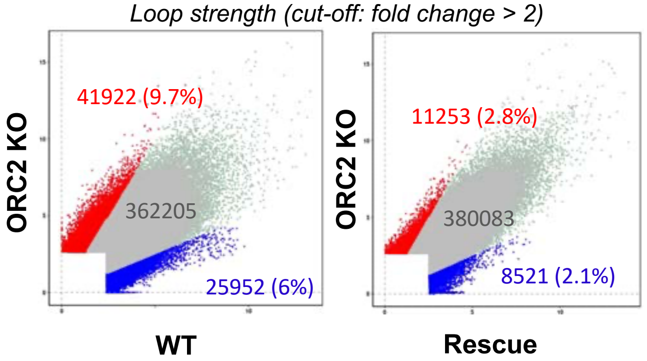


**F**

| **Z-score** | **Overlap w. Loop-up** |
|---|---|
| ATAC-up | 39.00 *** |
| ATAC-down | 7.70 *** |
| H3K9me3-up | -3.18 ** |
| H3K9me3-down | 0.39 |

**Figure S7.**

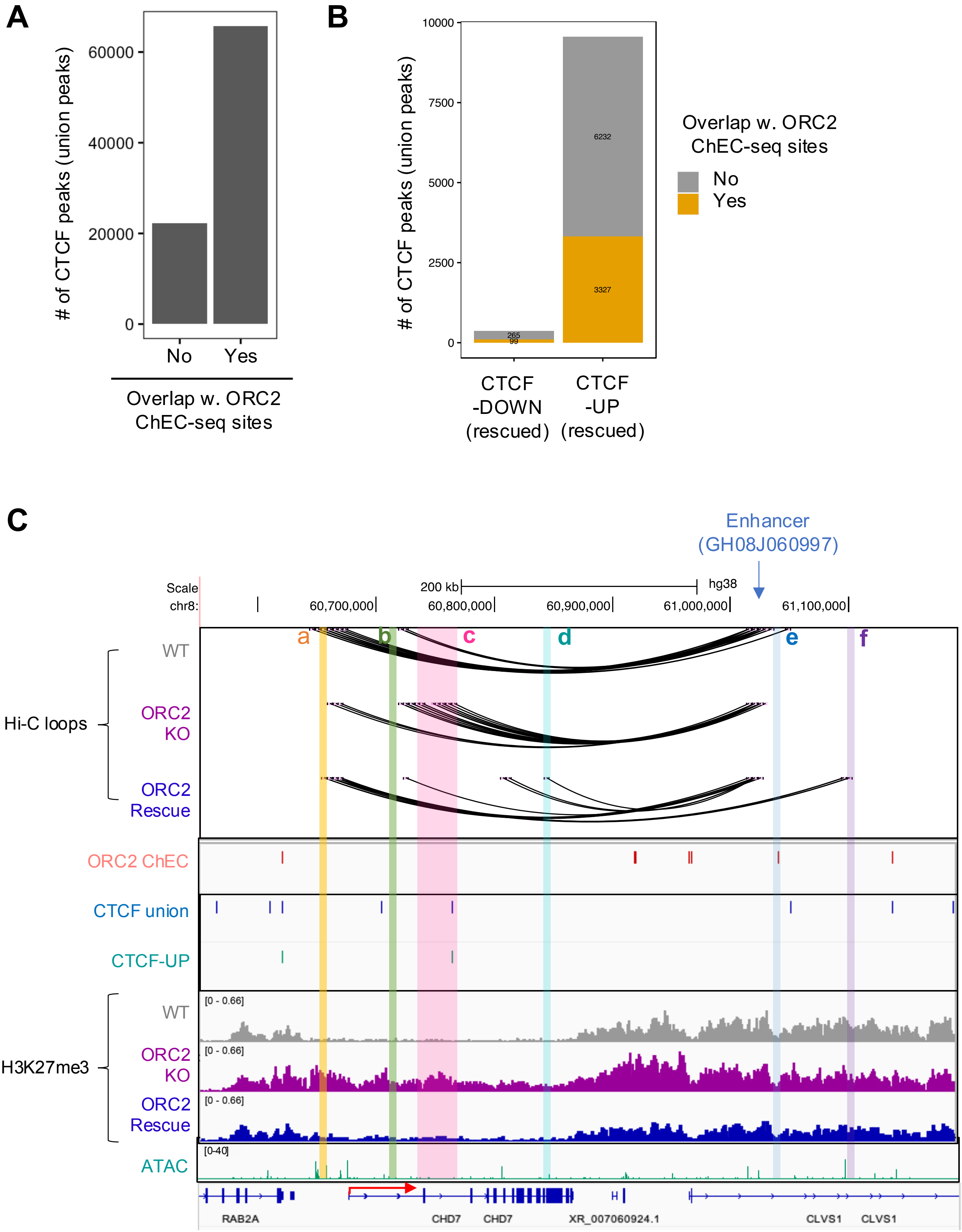
A
### of CTCF peaks (union peaks)
60000
40000
20000
0
No
Yes
Overlap w. ORC2 ChEC-seq sites
B
10000
7500
5000
2500
0
6232
3327
265
99
CTCF -DOWN (rescued)
CTCF -UP (rescued)
Overlap w. ORC2 ChEC-seq sites
No
Yes
C
Enhancer (GH08J060997)
Scale
chr8:
200 kb
hg38
60,700,000
60,800,000
60,900,000
61,000,000
61,100,000
a
b
c
d
e
f
Hi-C loops
WT
ORC2 KO
ORC2 Rescue
ORC2 ChEC
CTCF union
CTCF-UP
H3K27me3
[0 - 0.66]
ATAC
[0-40]
RAB2A
CHD7
CHD7
XR_007060924.1
CLVS1
CLVS1
